# vissE: A versatile tool to identify and visualise higher-order molecular phenotypes from functional enrichment analysis

**DOI:** 10.1101/2022.03.06.483195

**Authors:** Dharmesh D. Bhuva, Chin Wee Tan, Ning Liu, Holly J. Whitfield, Nicholas Papachristos, Sam Lee, Malvika Kharbanda, Ahmed Mohamed, Melissa J. Davis

## Abstract

Functional analysis of high throughput experiments using pathway analysis is now ubiquitous. Though powerful, these methods often produce thousands of redundant results owing to knowledgebase redundancies upstream. This scale of results hinders extensive exploration by biologists and often leads to investigator biases due to previous knowledge and expectations. To address this issue, we present vissE, a flexible network-based analysis method that summarises redundancies into biological themes and provides various analytical modules to characterise and visualise them with respect to the underlying data, thus providing a comprehensive view of the biological system. We demonstrate vissE’s versatility by applying it to three different technologies: bulk, single-cell and spatial transcriptomics. Applying vissE to a factor analysis of a breast cancer spatial transcriptomic data, we identified stromal phenotypes that support tumour dissemination. Its adaptability allows vissE to enhance all existing gene-set enrichment and pathway analysis workflows, removing investigator bias from molecular discovery.

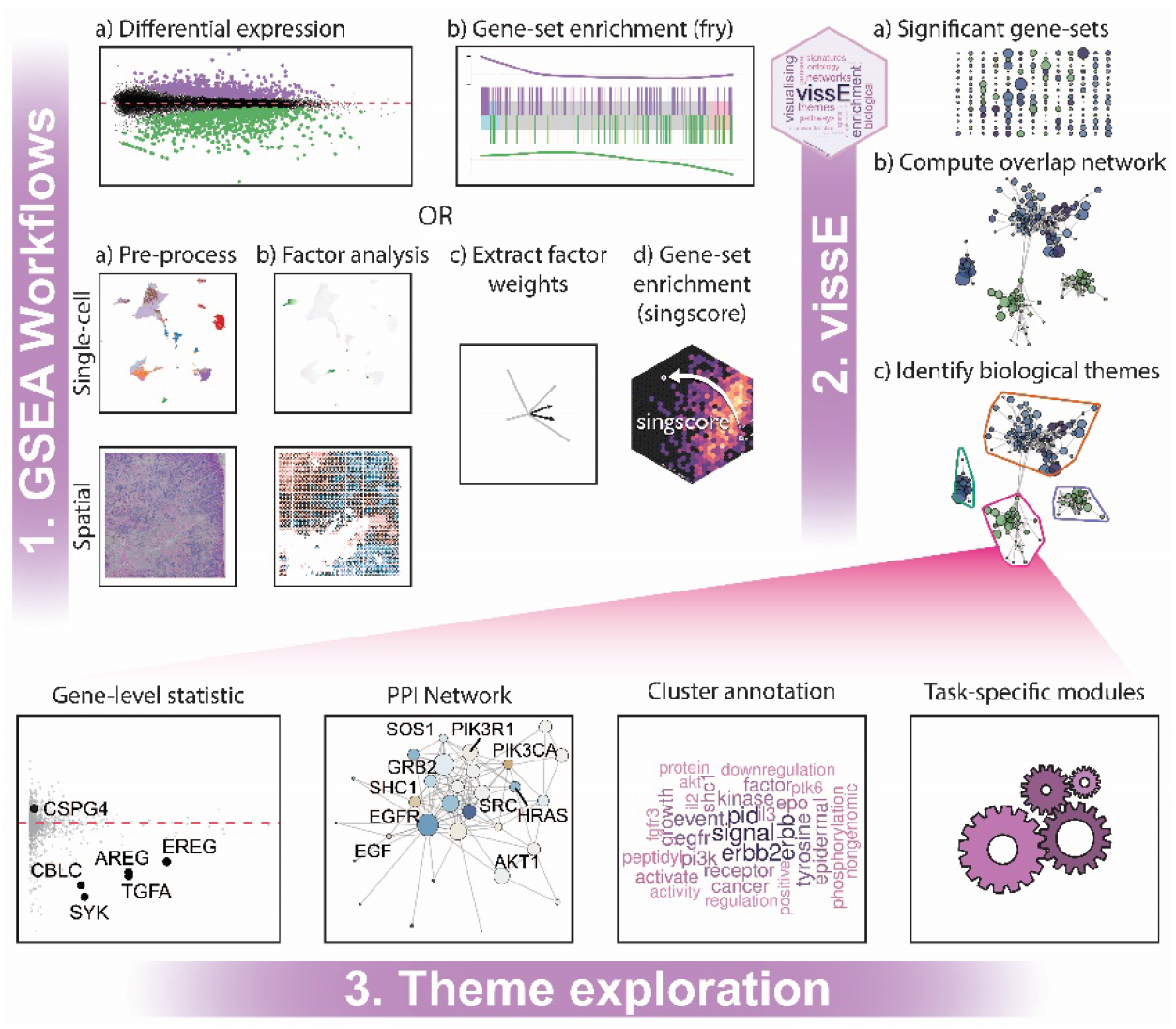

## Introduction

Biological systems are often studied using experiments that generate vast amounts of molecular measurements. Rigorous statistical analyses are routinely performed to identify the key molecules participating in the system. This is followed by interpretation from biologists who then attempt to explain the observed molecular shifts in their experiments, find evidence for molecular mechanisms and identify novel biology. Interpreting lists of molecules can be difficult and laborious for biologists in cases where thousands of molecules change in the experiment ^1^. This problem has motivated the development of statistical analyses that identify molecular processes enriched in the list of molecules, thus providing biologists with higher-order interpretable summaries of their experiments ^2^. When the molecules of interest are genes, these analyses have taken up the form of gene-set enrichment analysis ^2,3^ where sets of genes representing a common biological process are statistically assessed for enrichment in the experiment. Such gene-sets are derived from multiple sources which typically include the Gene Ontology (GO) project ^4^, the Reactome pathway database ^5^, the Kyoto Encyclopedia of Genes and Genomes (KEGG) pathway database ^6^ and the molecular signatures database (MSigDB) ^7,8^. These gene-sets are either curated from existing scientific literature or derived from molecular experiments.

Though gene-set enrichment analyses are powerful tools to study the biological processes underpinning biological systems, they often identify thousands of processes thus introducing a challenge in the interpretation of results. This is in part driven by redundancy introduced by the hierarchical structuring of processes in gene-set databases such as the GO, Reactome and KEGG ^9–11^. Additionally, the increasing number of experimentally derived gene-sets in databases such as the MSigDB will naturally lead to gene-set redundancy when related processes are being studied. Information redundancy in such databases is not necessarily detrimental, especially when evidenced by independent studies, however, it does pose a significant challenge when interpreting the results of gene-set enrichment analyses. Related pathways/processes are likely to be significant because of shared significant genes from the upstream analysis. In such a setting, biologists interpreting the top N processes will end up investigating the same signal in the data and will miss any orthogonal signals that although not as obvious, may lead to new insight into the nature of the data. Alternatively, domain experts attempting to interpret the full result set will inevitably be biased by their previous knowledge of the systems and will tend to select and focus on familiar processes for further investigations.

Three broad categories of solutions have been developed to address this problem: collapsing redundant information (for example, the creation of GO “slim” ontologies), incorporating redundancy information in the gene-set enrichment analysis method, or visualising redundancy in the gene-set enrichment analysis results. The first category focuses on modifying the underlying database such that redundant information is collapsed to produce a reduced collection of discrete categories, an approach that has been applied to GO and the Hallmark gene-sets from MSigDB ^7,12,13^. The second category of methods has been primarily developed for GO where the graph structure is incorporated into the statistical testing framework ^14–16^. While powerful, their application is limited to the analysis of knowledgebases that have a hierarchical structure such as GO and Reactome, and they can only be applied to a single database at a time restricting the sources of gene-sets.

On the other hand, the category of visualisation methods aims to reveal the redundancy structure in gene-set databases. As such, these methods can be coupled with a broader range of existing enrichment analysis methods ^2,3^, including some of the recent single-sample gene-set enrichment analysis methods ^17^. These methods generally begin by computing pairwise gene-set similarities based on content similarity of shared genes ^1,10,18–22^ or semantic similarity computed from the underlying graph structure ^23^. Gene-set clusters are subsequently identified by clustering on the similarity matrix directly ^11,18,22^ or by constructing a graph and applying graph clustering algorithms ^1,10,19–21^. The resulting similarity graphs are visualised with gene-set statistics such as p-values or enrichment scores overlayed onto vertices. A subset of methods attempt to collapse gene-set annotations in each cluster into a per-cluster annotation by either annotating each cluster using a single representative significant member ^11,18,21,22^ or by performing text-mining on all member gene-sets to identify an overarching biological theme ^1,10,20^. Table 1 summarises current methods developed to consolidate and visualise gene-set enrichment analysis results, detailing the approaches used to compute similarity, perform clustering, and annotate clusters.

**Table 1:** Methods to visualise the results of gene-set enrichment analyses

| Method | Similarity | Clustering | Annotation | Citation |
| --- | --- | --- | --- | --- |
| BiNGO | GO Structure | - | - | <sup>24</sup> |
| ClueGO | Cohen's Kappa | Similarity-based | Most significant term | <sup>18</sup> |
| EnrichmentMap | Jaccard index, Overlap coefficient | Graph (MCL) | Text-mining (word frequencies) | <sup>1,10</sup> |
| enrichplot | Semantic similarity, Jaccard index | - | - | 25,26 |
| GOMCL | Jaccard index, Overlap coefficient | Graph (MCL) | Most significant term | 21 |
| GOsummaries | - | - | Text-mining (weighted word frequencies) | 27 |
| GScluster | Overlap coefficient, Cohen's Kappa | Similarity-based (fuzzy clustering) | - | 28 |
| KOBAS-i | Jaccard index | Graph (Infomap) | - | 19 |
| Metascape | Cohen's Kappa | Similarity-based (hierarchical) | Most significant term | 22 |
| REVIGO | Semantic similarity | Similarity-based (hierarchical) | Most informative common ancestor | 11 |
| RICHNET | Jaccard index | Graph (edge betweenness) | Text-mining (word frequencies) | 20 |

Visualisation approaches are appealing since they can be applied to any gene-set enrichment analysis workflow. Good visualisations should reveal structure in the data that is not otherwise obvious. The approaches developed in the past have been powerful but restrict problem formulation to that of summarising the results of gene-set enrichment analysis, thus they lack the ability to provide useful insight from data. Furthermore, they lack suitable tools to visualise, explore and interpret the results of a gene-set enrichment analysis in the context of the underlying data and its downstream analysis. Tools such as EnrichmentMap ^10^ provide heatmaps to allow exploration of either the gene-level statistics (e.g., logFC) or their expression values, but not both. Other tools within the popular clusterProfiler software ^25,26^ allow exploration of gene-sets with respect to genes but fail to identify and characterise gene-set clusters. To address this limitation in problem formulation, vissE formulates the problem as one of identifying and characterising higher-order biological processes with the aim of allowing greater application and utility in all areas of biological and clinical sciences. Higher-order biological processes are identified by clustering on the gene-set network and are explored using various analytical modules including cluster annotation. Due to the open problem formulation, we can extend development of tools and are able to perform novel functional analysis workflows, such as unsupervised exploration of molecular phenotypes in single-cell and spatial transcriptomics data. The methods and data described here have been implemented in the `vissE`, `msigdb` and `emtdata` R/Bioconductor packages.

## Materials and methods

### A description of the vissE methodology

Gene-sets used in gene-set enrichment analysis often vary in the resolution of molecular phenotypes they represent. Different resolutions can therefore map onto the same biological process, and it is often of interest to identify the higher-order biological process that encapsulates related gene-sets. This idea is used in vissE to identify higher-order molecular phenotypes from a cluster of gene-sets of interest. Given a list of gene-sets, the pairwise gene-set similarity is first computed using either the Adjusted Rand Index (ARI), the Jaccard index or the overlap coefficient into a similarity matrix. When working with multiple databases, the Adjusted Rand Index or the Jaccard index are preferred since the overlap coefficient specifically highlights parent-child relationships and therefore works best when using a single hierarchically structured database. A gene-set overlap graph is generated by appropriately thresholding the similarity matrix. Under the assumption that gene-sets with many shared genes will likely represent related biological processes, vissE aims to identify clusters of gene-sets by applying graph clustering algorithms that harness topological information in the network. The preferred choice here is an algorithm based on random walks as this has been shown to work well for both dense and sparse graphs in identifying small and large community structures ^29^. Clusters are then ordered to maximise the statistic of interest, as well as their size, using the product of ranks approach described in ^30^.

The next step in a vissE analysis is to characterise each gene-set cluster and interpret the higher-order biological processes they represent using the range of analytical modules available in vissE. The vissE software provides a text-based interpretation of each gene-set cluster. This is done by performing text-mining analysis on the names or short descriptions of each gene-set in the cluster. The text data are pre-processed, and a term frequency is computed for each word. Concurrently, terms from the molecular signatures database (MSigDB) are pre-processed to compute inverse document frequencies (IDF) per term. Term frequencies within each gene-set cluster are then weighted against IDFs thus producing a term frequency inverse document frequency (TF-IDF) for each term. This helps to remove any database specific bias by down-weighting over-represented words in the database. To ensure the visualisation is not too dense, up to 25 words with the highest TF-IDF are used to represent each gene-set cluster and are visualised using word clouds as shown in Figure 1. These visual representations of biological themes summarise hundreds of enriched terms and are more conducive for interpretation by the user. Biologists can draw insights from these visualisations to interpret the biological processes represented by clusters of gene sets and quickly interpret gene-set enrichment analyses in the context of their data.

**Figure 1.**
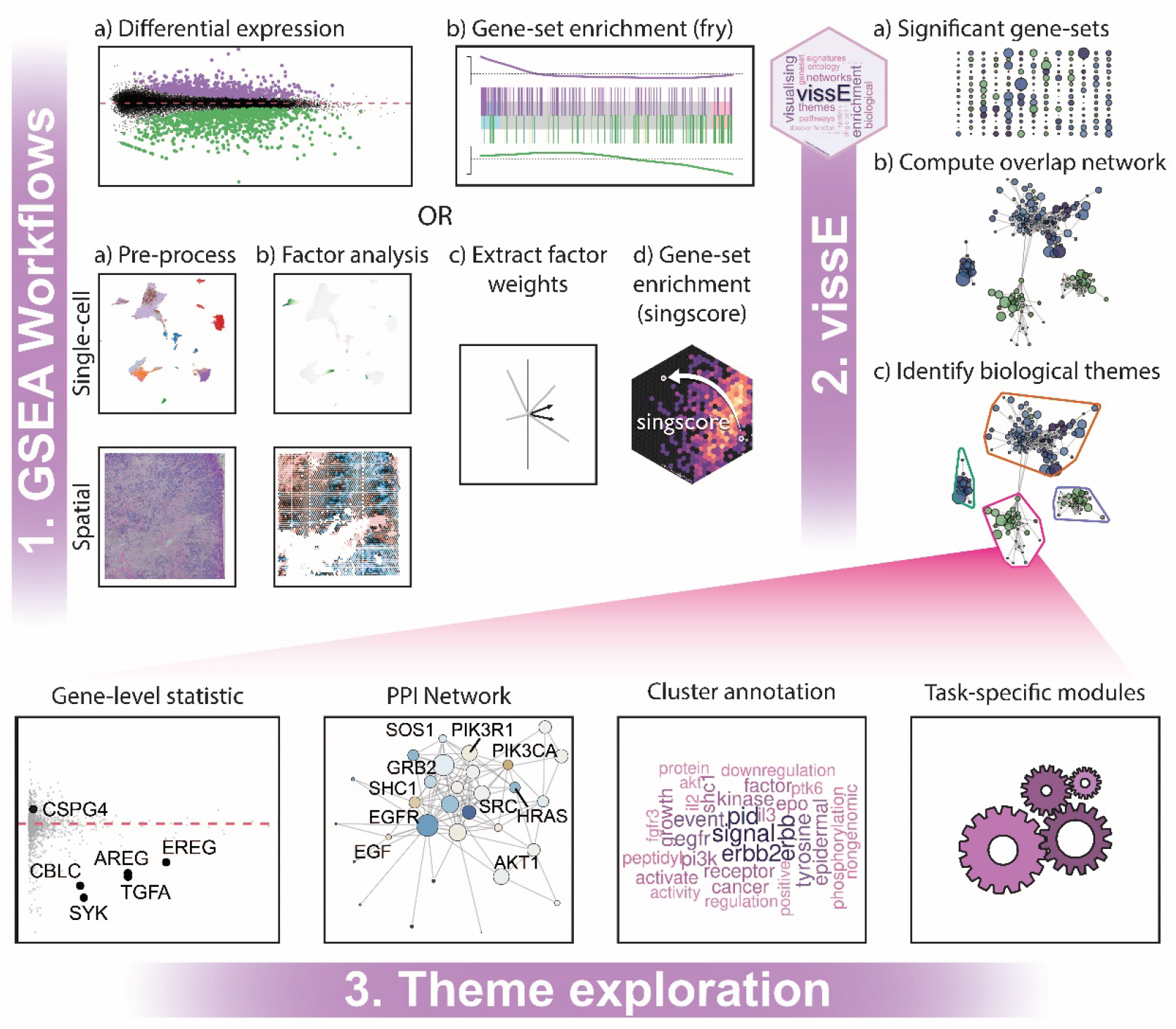
A schematic representation of various vissE workflows. A vissE workflow builds interpretable visualisations from gene-set enrichment analyses that allow users to easily investigate phenotypes at the resolution of biological themes and individual genes, while minimising investigator biases. (1) vissE is flexible for use with any gene-set enrichment analysis, including those from scRNA-sequencing, spatial transcriptomics and traditional bulk RNA-sequencing technologies. (2) A list of significant gene-sets from these analyses are used to generate a gene-set network that is used to minimise gene-set redundancy by identifying higher-order biological themes. (3) vissE offers a variety of analytical modules to then explore functional themes and to build a biological narrative that describes the underlying biological system being explored.

In addition to the word cloud, vissE enables characterisation of gene-set clusters using individual genes based on specific gene-level statistics of interest. In the context of a differential expression analysis, this could be the log fold change of each gene. A gene statistics plots comprising the statistic of interest of each gene and the number of gene-sets in the cluster that a gene belongs to can be generated (Figure 1). These gene statistics could be used to infer gene relevance within a process in the context of the specific experiment. High frequency genes could be interpreted as representative of clusters, and by extension, the associated higher-order biological process. Genes with both high statistics and frequencies within a cluster would be of interest in explaining the cluster with regards to the experiment. An analytical module within vissE also allows visualisation of protein-protein interactions between genes in a gene-set cluster, providing an independent line of evidence for common underlying higher-order biological processes. These analytical modules collectively allow vissE to be a powerful tool for understanding the higher-order processes identified as show in Figure 1.

### Deriving the overlap graph

Pairwise gene-set similarities for gene-sets of interest, such as those that are significant in a gene-set enrichment analysis, are computed using the Adjusted Rand Index (ARI), the Jaccard Index or the overlap coefficient. A gene-set overlap graph is built by thresholding the similarity matrix and represented using the `igraph` R package ^31^. Gene-sets without any connections are dropped from the graph. All graphs in the package are visualised using the `ggraph` R package.

### Pre-processing text for text-mining analysis

Text used to annotate gene-set clusters is first split using the “_”, “/”, “@”, “(”, “)” and “|” characters to generate words. Punctuation marks, extra white spaces, words that are numbers, stop words used in the English language and user provided blacklisted words are removed. Words such as KEGG, hallmark and other such prefixes and suffixes commonly used to name or describe gene-sets are removed by default. All characters are transformed to lowercase characters. String lemmatisation is performed to produce lemmatised words. All text-mining analysis is performed using the `tm` R package ^32^.

### Developing a protein-protein interaction network

The international molecular exchange consortium data was used to produce a protein-protein interaction (PPI) network for human and mouse ^33^. The full PPI was downloaded in the PSI-MI TAB format (as of 6^th^ July 2021) and records where both the source and target nodes are of the same species were retained. The human and mouse PPIs were then filtered out and separated. Uniprot IDs were mapped to Entrez IDs to allow gene-level queries from vissE. Human to mouse ortholog data from the HGNC Comparison of Orthology Predictions (HCOP) database ^34^ were used to infer PPIs for each organism using the other. This was done to provide better coverage for organisms like mouse that have not been studied as extensively. Inferred interactions were annotated in the resulting data and can therefore be filtered out when necessary. Duplicated edges, defined as interactions involving the same two Entrez IDs, were combined with the maximum confidence score taken as the confidence score. The resulting PPIs are available from the `msigdb` R/Bioconductor package.

### Processing bulk RNA-seq data

Sequencing reads from ^35^ were downloaded from the sequencing reads archive (SRA) using `sratools`. The Subread ^36^ aligner was used to align reads to the GRCh38 human reference genome and `featureCounts` ^37^ was used to quantify reads per gene. Genes with low expression were filtered out and normalisation factors were calculated using the TMM method ^38^. This data is made available through the `emtdata` R/Bioconductor package. Differential expression analysis was performed using the quasi-likelihood pipeline ^39^ from the `edgeR` R/Bioconductor package ^40^. Gene-sets from the hallmark collection (h), Reactome, KEGG, WikiPathways, and the gene ontology were downloaded from the molecular signatures database (MSigDB v7.2) using the `msigdb` R/Bioconductor package. These were then used to perform gene-set enrichment analysis using the limma::fry method.

### Processing scRNA-seq data

Pre-processed data from ^41^ was downloaded from the gene expression omnibus (GSE161529) with quality control performed as described in the original publication. Three ER+ breast cancer samples (IDs), three BRCA1 triple negative breast cancers (IDs) and one triple negative breast cancer sample (ID) were selected for further analysis. Data from each sample were normalised using the `scran` R package ^42^ and then integrated by identifying integration anchors using the Seurat R package ^43^. PCA was performed on the top 2000 highly variable genes defined based on the mean-variance relationship of genes. Cell type annotation was performed using the `SingleR` ^44^ and `scClassify` ^45^ R packages. Cells were first annotated using the human primary cell atlas data (HPCA) ^46^ and ovarian cancer data ^47^ independently using SingleR. Endothelial cells identified using the HPCA reference were annotated as such. From the remainder of cells, malignant cells and fibroblasts were annotated using the ovarian cancer data as a reference. The remaining cells were annotated using the HPCA as a reference. T-cells identified using the HPCA as a reference were further annotated using the joint estimation model of scClassify to further sub-divide T-cell subtypes. Malignant cells were further annotated as estrogen receptor positive (ER+), triple negative breast cancer (TNBC) or TNBC BRCA1-mutant based on the subtype of the patient they originated from. Data were visualised using uniform manifold approximations (UMAPs) computed from the first 50 principal components (PCs) using implementations in the `scater` R package.

### Processing spatial transcriptomics data

Visium spatial targeted data of human invasive lobular carcinoma breast tissue (ER positive, PR positive, HER2 negative) used in this study was obtained from the 10X Genomics demonstration datasets ^48^. Data were pre-processed using Space Ranger software v1.2.0. Spots with library sizes smaller than 3000 and less than 500 expressed genes were filtered out while the rest were normalised using the `scran` R package ^42^. PCA was performed on the top 2000 highly variable genes defined based on the mean-variance relationship of genes. Cell type deconvolution was performed with the RCTD method ^49^ using a single-cell dataset (GSM4909302) from the previous section as a reference.

Spots mapping stroma surrounded by different types of malignant cells were defined by mapping the pathologist’s annotations onto the spatial transcriptomics data. Pixels within a 150-pixel circular radius were used to define spots. Spots with more than 75% stromal annotated pixels were defined as stromal spots. The surroundings of stromal spots were defined based on a square grid. Windows starting at x-coordinates 3000 and 9000 pixels and of width 6000 pixels, and y-coordinates 16000 onwards, were defined as stroma surrounded by malignant mesenchymal cells and stroma surrounded by malignant epithelial cells respectively. Only the stromal spots within these windows were used for the differential expression analysis. Pseudo-replicates were defined by splitting windows within each group into three equally sized bins along the x-axis. Pseudo-bulk samples were subsequently created and subjected to a differential expression analysis ^39^ followed by a fry analysis, and finally a vissE analysis.

### Gene-set enrichment analysis of factors

Factors identified in a factor analysis often have loadings, amplitudes or weights representing feature importance. Principal components analysis (PCA) of RNA sequencing data produces gene loadings that reflect the relevance of each gene to the principal component (PC) of interest. Gene loadings can be used to compute gene-set scores that reflect the importance of each gene in the PC. We used the singscore method ^17^ implemented in the `singscore` R/Bioconductor package to compute gene-set scores for each gene-set in a PC for the single-cell and spatial transcriptomics datasets. In each PC, genes were ranked using their gene loadings. Scores were computed for all gene-sets in the hallmark collection (h), Reactome, KEGG, WikiPathways, gene ontology and the single-cell gene-sets collection (c8) of the molecular signatures database (MSigDB v7.2). This produced gene-set scores for these gene-sets in each PC identified using PCA.

### Running EnrichmentMap

The EnrichmentMap plugin (v3.3.2) in Cytoscape (v3.9.0) was used to identify and characterise higher-order phenotypes in the bulk RNA-seq data. Genes ranked based on logFCs were used to perform gene-set enrichment analysis using the GSEA method ^8^ as per the EnrichmentMap workflow ^1^. The gene-set database used was the same as that used for the vissE analysis. Default setting were used to generate the gene-set overlap graph, identify clusters and annotate clusters.

## Results

### Higher-order molecular phenotypes involved in an epithelial to mesenchymal transition in breast cancer

Here we demonstrate the application of vissE to a standard differential expression analysis. In epithelial tumours, malignant cells can undergo an epithelial to mesenchymal transition (EMT) and acquire mesenchymal properties such as migration and motility. The process of EMT is thought to enable cancers to metastasise ^50^. While often characterised as a single process, the transition from an epithelial to mesenchymal phenotype involves various complex changes to cells and their microenvironment ^51^. To explore these processes, we used data from the human mammary epithelial (HMLE) cell line system in Cursons, et al. ^35^ where a mesenchymal subline of the HMLE cell line (mesHMLE) was induced by TGFβ stimulation and maintained with epidermal growth factor (EGF). Differential expression analysis was performed followed by gene-set enrichment analysis that identified 1240 significant gene-sets at the FDR level 0.1. These gene-set were then processed using vissE to identify higher-order biological processes. A threshold of 0.25 was applied on the adjusted Rand index (ARI) to produce the gene-set overlap network. Disconnected gene-sets were dropped producing a network with 1170 nodes and 4113 edges. Community detection using the walktrap algorithm identified 195 non-overlapping gene-set clusters that were then characterised using tools within vissE. Figure 2 shows four higher-order processes that are expected to change during EMT, demonstrating how vissE captures key biological properties of a dataset.

**Figure 2.**
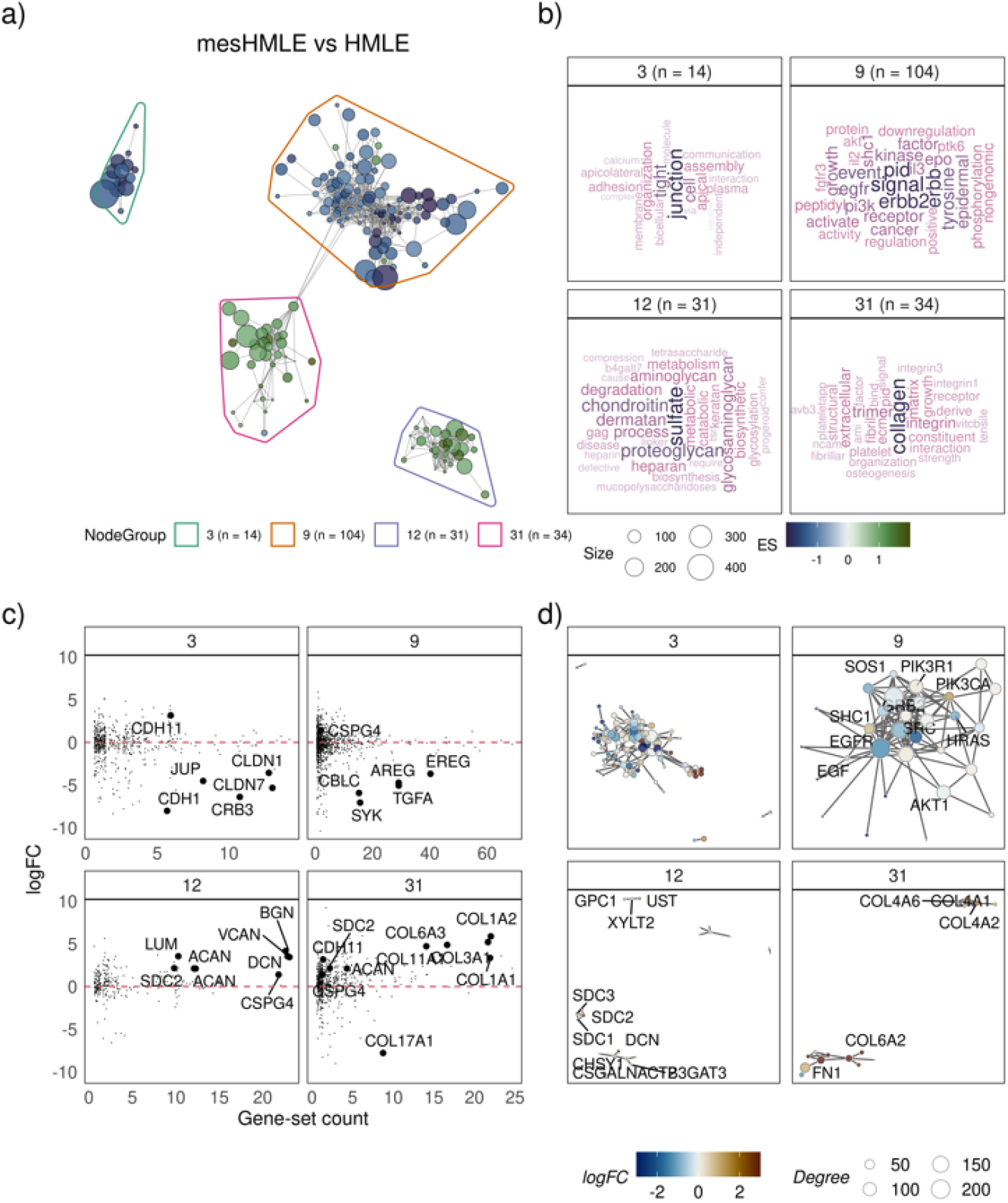
Using vissE to identify and characterise biological themes observed in an epithelial to mesenchymal transition (EMT) in the human mammary epithelial (HMLE) cell line. a) A gene-set overlap graph of gene-sets enriched during an EMT with nodes representing individual gene-sets and edges representing overlaps based on the adjusted rand index (ARI). Nodes are coloured based on the direction and significance of enrichment: green nodes represent gene-sets enriched in mesenchymal cells and blue in epithelial cells. Four gene-set clusters representing biological themes are identified, containing 14, 104, 31 and 34 gene-sets respectively. b) Cluster annotations generated by text-mining analysis of gene-set names. c) Log fold-change (logFC) of genes belonging to gene-sets in the cluster plot against the number of gene-sets in the cluster the gene belongs to. d) Protein-protein interaction (PPI) networks between genes that belong to gene-sets in the cluster. Each node represents a gene and edges represent known PPIs. Nodes are coloured based on the logFC.

We identified higher-order phenotypic changes associated with cell-cell interaction reflect the loss of junctions and cell-cell adhesion in epithelial cells that is necessary for their transition into a mesenchymal phenotype ^51^. Specifically, Figure 2 shows that cluster 3, identified by vissE, represents tight junctions that were downregulated in mesenchymal cells relative to the epithelial HMLE cells. This and other themes are recognised and interpretable when word clouds are generated either using set names or short descriptions (Additional File 1: Supplementary Figure 1). Key genes identified include claudin genes (*CLDNs*) such as *CLDN7* and E-cadherin (*CDH1*) (Figure 2c, cluster 3), that are known epithelial markers and are predictive of an epithelial state ^52^. Other than downstream phenotypic changes, vissE also captured changes in signalling such as differences in EGFR/HER2 signalling between HMLE and mesHMLE cell lines (Figure 2, cluster 9). Specifically following through the analysis of cluster 9 in Figure 2, EGFR/HER2 signalling (text as ‘erbb2 signal’) was relatively lower in mesHMLE compared to HMLE (Figure 2a) and protein interactions amongst key ERBB signalling proteins including ERBB2/3/4, EGFR and downstream signalling proteins like SHC1 and SOS2 were observed. The HMLE cell line has been demonstrated to depend on autocrine EGFR signalling for growth and proliferation ^53,54^, hence, it is expected that EGFR/HER2 signalling activity in HMLE is higher than in mesHMLE. Additionally, TGFβ transactivates EGFR in breast cancer ^55^ therefore removal of TGFβ stimulation in the mesHMLE subline attenuated EGFR/HER2 signalling as evidenced by the relative downregulation of EGF signalling ligands such as *AREG* in Figure 2c. However, since EGFR/HER2 signalling was not completely lost in the mesHMLE subline, its mesenchymal phenotype was stably maintained (AveLogCPM of *AREG* in mesHMLE was 4.321). The signalling events identified in cluster 9 clearly reflect the biology expected in this experiment and validates the vissE workflow. All other themes identified by vissE are included in Additional File 2.

Other than the known or expected processes, vissE was able to identify other processes of interests. Cluster 12 identified an up-regulated higher-order process involving proteoglycans such as *VCAN* and *SDC1*, and sulfate proteoglycans (*SPGs*). These genes are known to regulate cell adhesion and motility ^56^ and were mostly up regulated in mesenchymal HMLE cells as seen in Figure 2c. Similarly, cluster 31 represents numerous collagen genes that were up regulated in mesHMLE cells. Both these clusters represent different components of the extra cellular matrix (ECM). TGFβ signalling in mesenchymal cells has been known to directly affect accumulation of fibrillar collagens in the ECM ^57^ and the results from cluster 31 suggest this is also the case in the TGFβ stimulated HMLE system. Up-regulation of genes in clusters 12 and 31 suggest a more rigid ECM that promotes EMT via nuclear localisation of *TWIST1* ^58^. Our differential expression analysis supported this hypothesised mechanism as evidenced by the up regulation of TWIST1 in mesHMLE (logFC = 2.403, FDR = 0.002). Further validations of the mechanism that promote TGFβ-induced EMT is the up-regulation of proteoglycans in response to the growth factor, such as versican (*VCAN) ^59^* and aggrecan (*ACAN) ^60^*, which provides a favourable ECM for migrating mesenchymal cells and enables detachment of cells from the basement membrane ^61^. Collectively, the vissE analysis was able to identify and visualise these higher-order processes, capturing the cell-extracellular matrix remodelling that is required for EMT in a clear and unbiased manner.

We contrast vissE with two alternative analysis strategies common in the literature. In the first, we focus on the top N gene-sets from an enrichment analysis and in the second we compare to results from the EnrichmentMap tool. We assessed redundancy in the selected top N gene-sets by computing the degree of overlap of DE genes in the top 50 significant gene-sets. Many of the top 50 gene-sets shared a large number of DE genes, suggesting that their significance was attained due to the same set of underlying DE genes. Additionally, these gene-sets formed clusters based on their DE gene overlap demonstrating that the same sets of processes were captured repeatedly in the top 50 gene-sets (Additional File 1: Supplementary Figure 2). EnrichmentMap revealed many of the same processes we identified using vissE, however, the method clustered considerably fewer gene-sets for the biological themes it identified, for example, it identified only 6 gene-sets in the sulfate proteoglycan cluster as opposed to the 31 vissE identified (Additional File 1: Supplementary Figure 3). This was the case for most other biological themes, demonstrating that vissE provided better coverage of the enrichment results than EnrichmentMap. In some cases, the default cluster annotation from EnrichmentMap produced uninformative cluster labels such as labelling a cluster representing positive regulation of alpha and beta T-cell activity as “positive beta alpha”.

### De-novo identification of higher-order molecular phenotypes in single-cell RNA-seq experiments

Single-cell RNA-sequencing experiments are now commonly used to probe phenotypes associated with cell identity; molecular measurements at the cellular level can allow finer dissection of molecular phenotypes in a biological system. Powerful exploratory analysis without any presumptions on the biology can be performed with such high-resolution data. Unlike the bulk RNA-seq setting where we begin with a specific research question or hypothesis, such as a comparison between known groups, here we introduce a more flexible framework to explore molecular phenotypes. Very few approaches exist for this type of analysis of single-cell transcriptomic data. Firstly, factor analysis of the high-dimensional data is performed to identify factors that represent the underlying biological processes. In most cases, methods such as principal components analysis (PCA) are used to identify orthogonal factors that, in essence, reflect orthogonal groups of biological processes. Here, we used principal components analysis to identify the top 5 factors from a single-cell RNA-seq breast cancer dataset containing 51660 cells from seven patients across two breast cancer subtypes (Figure 3a). The identified factors were interpreted by performing gene-set enrichment analysis on each factor using singscore ^17^ as described in the methods. Higher-order phenotypes were then identified in each factor by performing a vissE analysis on gene-sets with absolute scores greater than 0.2. A Jaccard index threshold of 0.25 was applied in vissE to generate the gene-set overlap network.

**Figure 3.**
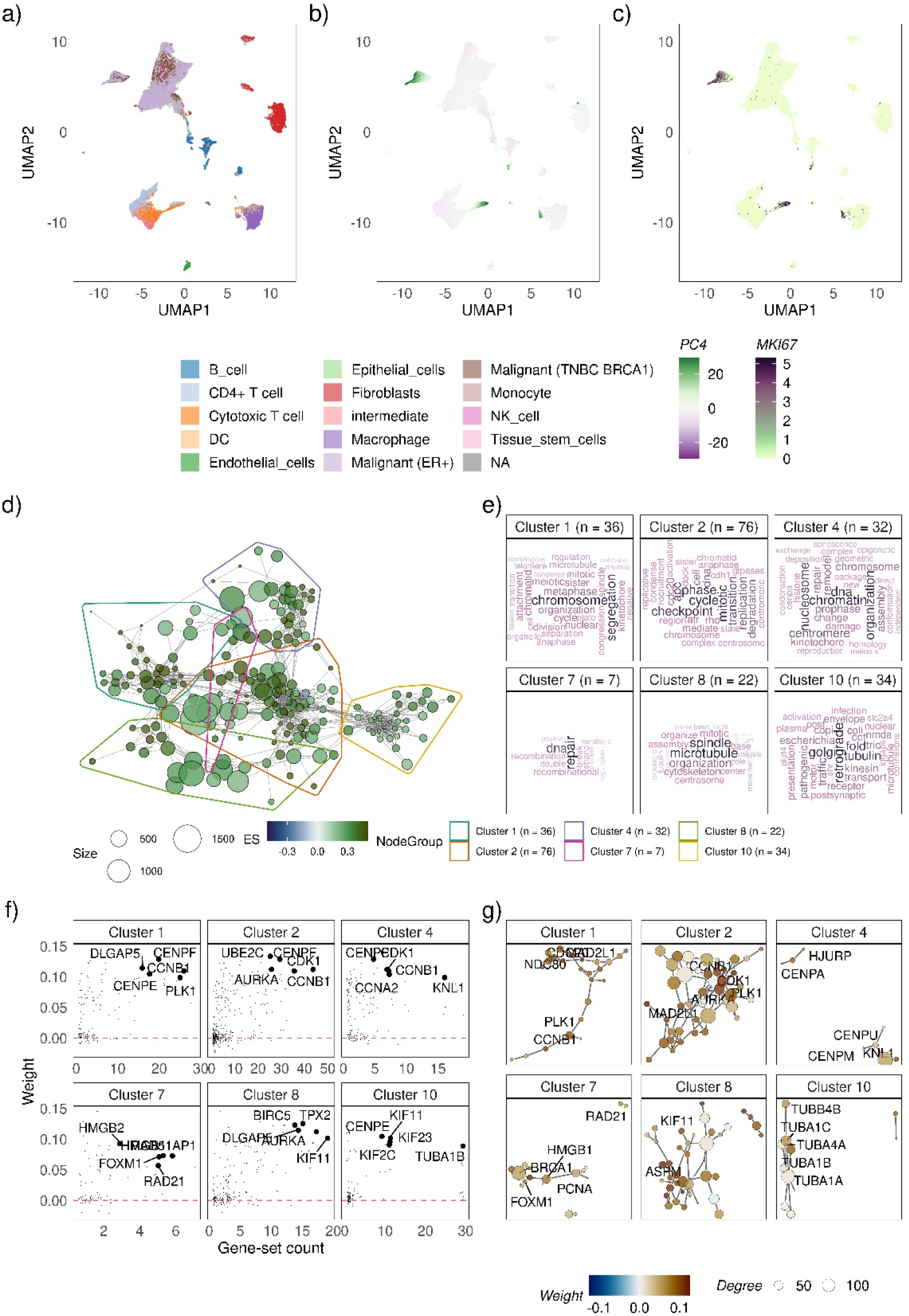
Using vissE to identify and characterise a proliferative phenotype in single-cell transcriptomic data of seven breast cancer patients from. a-c) A uniform manifold approximation projection (UMAP) of cells from 7 patients annotated by a) inferred cell types. b) the projection of the fourth principal component (PC4). c) expression of the MKI67 gene that encodes the Ki67 marker of proliferation. d) A gene-set overlap graph of gene-sets enriched in PC4 with nodes representing individual gene-sets and edges representing overlaps based on the adjusted rand index (ARI). Nodes are coloured based on the direction and significance of enrichment: green nodes represent gene-sets enriched in PC4 high cells. Six gene-set clusters representing biological themes are identified, containing 36, 76, 32, 7, 22 and 34 gene-sets respectively. e) Cluster annotations generated by text-mining analysis of gene-set names. f) Gene loadings (also known as weights) for genes belonging to gene-sets in the cluster plot against the number of gene-sets in the cluster the gene belongs to. g) Protein-protein interaction (PPI) networks between genes that belong to gene-sets in the cluster. Each node represents a gene and edges represent known PPIs. Nodes are coloured based on gene loadings (also known as weight).

Figure 3 shows the results when vissE and singscore are applied to the fourth principal component (PC4) of the data. Figures 3a-b show the UMAP projections of cells with the cell type and the PCA projection on factor 4 (i.e., PC4) annotated accordingly. These plots show that the molecular phenotype identified by the fourth principal component does not represent a cell type nor cells from a single patient (Additional File 1: Supplementary Figure 4) but a phenotype that is common to cells from various cell types, including malignant cells and immune cells. Functional analysis of Factor 4 (PC4) using singscore identified 704 gene-sets with absolute scores greater than 0.2. Analysing these gene-set using vissE with an ARI threshold of 0.3 identified 82 gene-set clusters. Six representative clusters shown in Figures 3d-g clearly reveal a proliferative phenotype that is present in a subset of cancer and immune cells (as shown in panel a-c, other clusters found in Additional File 3).

Specifically, cluster 2, the largest cluster in Figure 3d represents the broader set of gene-sets associated with the cell cycle as evident from the word cloud in Figure 3e and the genes highlighted in Figure 3f. These clusters capture gene-sets related to cell proliferation, including cell cycle stages (cluster 2) or organelle activity such as chromosome segregation (cluster 1), centrosomic changes (cluster 4) and microtubule formation (cluster 8). Most genes in these representative clusters have positive PC loadings (as shown by the gene-level statistics/weights in Figure 3g) suggesting they are positively associated with factor 4 and by extent, the process of proliferation. Clusters 7 and 10 represent processes that are required for a smooth transition through the cell cycle. DNA damage repair is required to ensure error free replication ^62^ and the secretome pathway of retrograde transport via the Golgi is required for recycling membrane bound proteins during cell division ^63^. These clusters are themselves heavily interlinked indicating a strong dependence between the processes they represent.

The gene-level statistics in Figure 3g can link interpretations back to specific genes, enabling the identification of key regulators or markers of the processes identified. For instance, genes identified in clusters 1, 2, 4 and 8 such as *AURKA* and *CDK1* are known kinases regulating cell cycle progression ^64^. Furthermore, vissE also provides the protein-protein interaction network (Figure 3f) that serve as a line of evidence independent from the enrichment analysis and/or gene set membership that can be further explored using specialised network analysis tools to identify key proteins in the relevant processes. Collectively, these findings suggest that factor 4 is identifying a subpopulation of proliferating cells as validated by the expression of the *MKI67* gene (Figure 3c). They also showcase how vissE captures shared phenotypic characteristics that span several cell types across various patients.

### Higher-order spatially resolved molecular phenotypes of tumour promoting cancer associated fibroblasts

The advent of spatially resolved transcriptomics data has enhanced the context-specific exploration of biology. The factor analysis pipeline described in the previous section can be used to perform an unbiased exploration of molecular phenotypes in any transcriptomic data, including spatial transcriptomics data. We applied the factor analysis pipeline to a human invasive lobular carcinoma breast tissue (estrogen receptor positive, progesterone receptor positive, and HER2 negative) dataset ^48^ that contains transcriptomics measurements profiled across 4325 spots. The data were pre-processed, and factors were identified by applying PCA to the 3364 spots that passed quality control (see Methods). The top 5 factors identified were subjected to a gene-set enrichment analysis using singscore ^17^ and resulting gene-sets with absolute scores greater than 0.2 were interpreted using vissE by applying an ARI threshold of 0.2.

The original H&E stained tissue slide (Figure 4a) was profiled using spatial transcriptomics and annotated by a pathologist for regions of stroma, malignant epithelial cells, and malignant mesenchymal cells (Figure 4b). Spots were projected onto PC1 and subsequently mapped onto the original spatial landscape to explore and characterise the resultant spatial patterns (Figure 4c). A key finding was that the gene expression pattern of stroma adjacent to epithelial cells differed from the stroma adjacent to mesenchymal-like cells (Figure 4b). Regions with positive PC1 projections captured stroma infiltrated by mesenchymal-like malignant cells (Figure 4b-c). The singscore analysis of this PC identified 880 gene sets that were then clustered into 107 biological themes using vissE. Our vissE analysis showed that these regions were enriched in collagen-related (cluster 1), sulfate proteoglycan metabolism (cluster 4) and other cell-ECM binding (cluster 13) gene-sets, characterising the tumour-stromal interactions between cell populations at the boundaries of the tumour (Figure 4d-g, other clusters found in Additional File 4). Positive gene weights (Figure 4f) for stroma-specific genes, including collagens (e.g., *COL4A1*), *VCAN*, *FN1* and many ECM proteins (Figure 4f-g), further indicated ECM remodelling and suggested the contribution of fibroblasts to this transcriptomic signal. Gene-sets relating to growth factor expression (cluster 11) and chemotaxis (clusters 11, 12 and 25) were also enriched. In addition, cluster 25 relates to the regulation of VEGF-induced migration, including the expression of key VEGF-related genes (*NRP1, NRP2, FLT1, KDR, PGF*), which can promote tumour dissemination by supporting the invasion of malignant cells into the stroma. These results, coupled with the up-regulation of chemokines (*CXCL12*) and growth factors (PDGF and TGFβ, see Figure 4g) reflect the tumour-stromal interactions in this tumour microenvironment that support the invasion of mesenchymal-like malignant cells in adjacent stroma.

**Figure 4.**
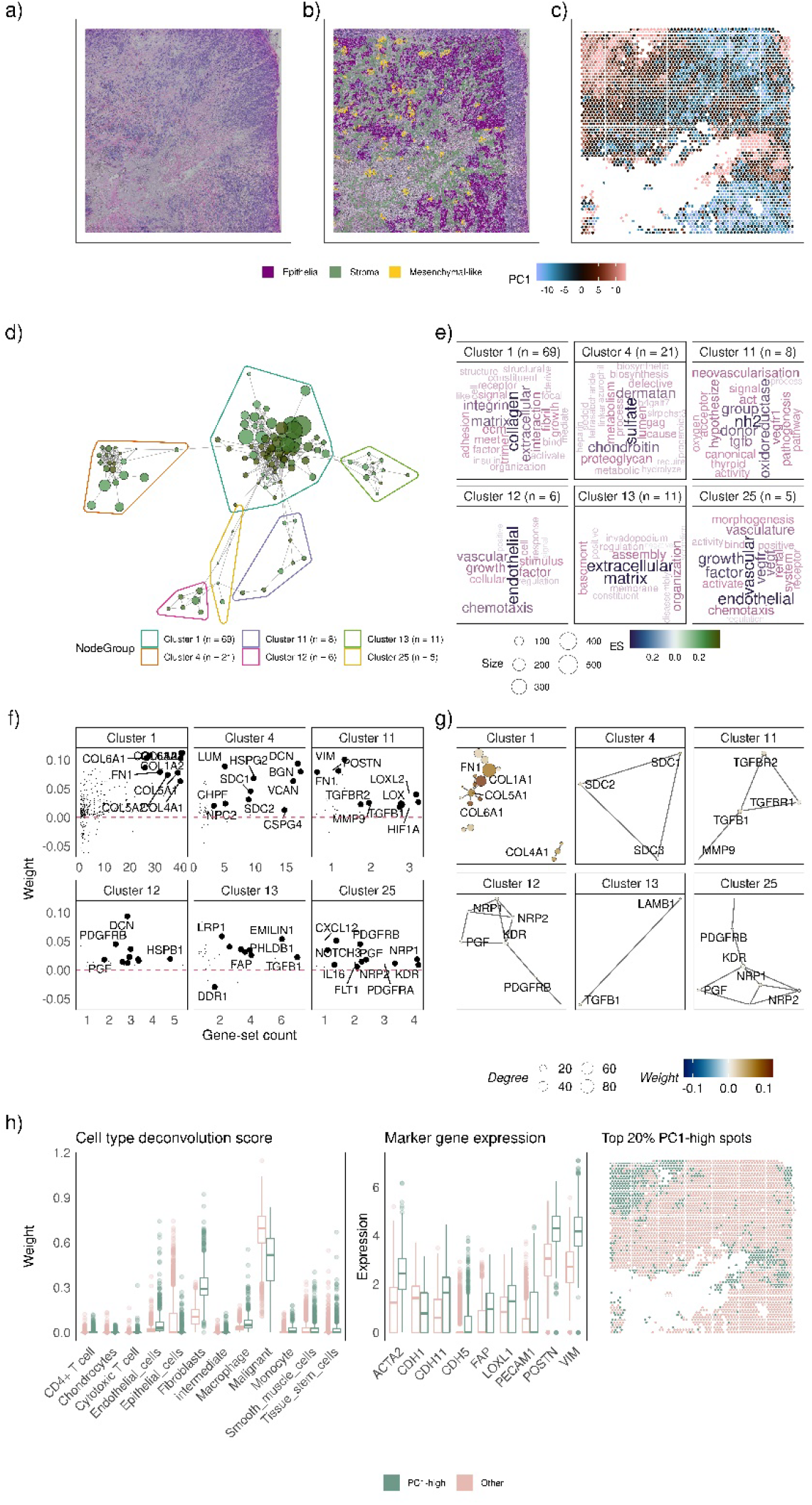
Using vissE to identify and characterise a cancer associated fibroblast (CAF) phenotype in spatial transcriptomics data of a breast cancer patient. a) A H&E image of the breast cancer tissue profiled using the 10X visium technology. b) Spots profiled coloured by the projection of the first principal component (PC4). c) Pathologist’s annotations of stromal (olive-green), malignant (purple) and mesenchymal-like (gold) regions of the tissue overlayed on the H&E image. d) A gene-set overlap graph of gene-sets enriched in PC1 with nodes representing individual gene-sets and edges representing overlaps based on the adjusted rand index (ARI). Nodes are coloured based on the direction and significance of enrichment: green nodes represent gene-sets enriched in PC1-high spots. Six gene-set clusters representing biological themes are identified, containing 69, 21, 8, 6, 11 and 5 gene-sets respectively. e) Cluster annotations generated by text-mining analysis of gene-set names. f) Gene loadings (also known as weights) for genes belonging to gene-sets in the cluster plot against the number of gene-sets in the cluster the gene belongs to. g) Protein-protein interaction (PPI) networks between genes that belong to gene-sets in the cluster. Each node represents a gene and edges represent known PPIs. Nodes are coloured based on gene loadings (also known as weight). h) Cell type deconvolution (left) and expression of CAF-related marker genes (center) for the top 20% of spots with the highest PC1 projection vs. all other spots (region marked in the right panel).

These higher-order biological themes point to a cancer-associated fibroblast (CAF) phenotype. Biomarkers of CAFs like *PDGFRA*, *PDGFRB*, *TGFB1* (cluster 12 and 25), *FAP* (cluster 13), *MMP9* (cluster 11), *LOXL1* and αSMA (also known as *ACTA2*) are more highly expressed in PC1-high regions (top 20% spots) as seen in Figure 4h. Cell type deconvolution results agreed with this hypothesis. PC1-high regions demonstrated strong evidence for fibroblasts and weak support for malignant cells (Figure 4h). The few malignant cells present in these regions (Figure 4h) could contribute to CAF formation via TGFβ signalling as evidenced by the upregulation of TGFβ1 and its receptors (cluster 11) ^65^. ECM stiffening induces mechanical stress that further activates CAFs ^66^. Upregulation of the CAF-induced pre-metastatic niche (PMN) marker *POSTN* (cluster 11) ^67^, chemotaxis and vasculature (clusters 12 and 25) as well as higher deconvolution weights of endothelial cells and macrophages in the PC1-high regions are evidence for a tumour promoting role of CAFs at the leading edge of tumours ^68^.

To validate these findings, we performed a supervised differential expression analysis of the stroma surrounded by different types of malignant cells defined using our pathologist’s annotations. We found that gene expression signatures and higher-order themes identified in our unsupervised PCA analysis (Figure 4) were consistent with those identified in our supervised analysis of the stroma (Additional File 1: Supplementary Figure 5), demonstrating that these stromal regions are secreting extracellular matrix constituents and remodelling the ECM to support the invasion of mesenchymal cancer cells.

## Discussion

Functional interpretation of high-dimensional molecular data has been a challenge since the advent of high through-put technologies. The rate of data generation greatly outcompetes the rate of their analysis and interpretation, leaving many data under-explored. While statistical and computational tools have assisted in identifying molecules/features of interest in data, these results are difficult to interpret functionally. Gene-set enrichment analysis is a solution to functional exploration of molecular data; however, it results in the identification of numerous biological processes and often limits a holistic interpretation of the data. In such scenarios, it is common to use the top significant processes to understand the biological system being studied. In this study, we showed that while such an approach will control the FDR at a desired level, the top gene-sets would provide redundant biological insight because of a shared set of significant genes/molecules (Additional File 1: Supplementary Figure 2). This effect would be amplified when hierarchically structured knowledgebases are used. The vissE method tackles gene-set redundancy by condensing information from all significant gene-sets into higher-order biological processes, thus hierarchically structuring the results in an easily browsable manner: starting with identification of higher-order processes of interest, then dissecting the gene-sets within that process, and finally drilling down to the genes common across those gene-sets. Associations between different higher-order processes can also be explored providing a more comprehensive landscape of the system being studied.

The redundancy of biological knowledge both within and between knowledgebases is exploited by vissE to enable robust identification of higher-order processes. Within-knowledgebase redundancy helps derive higher-order processes while between-knowledgebase redundancy provides additional independent evidence of processes. As such, vissE can accumulate and structure functional evidence derived from gene-set enrichment analysis methods. Accumulation of gene-sets across knowledgebases can also assist in reducing the impact of poor-quality gene-sets as their effect would be averaged out. A caveat to collecting information from across sources is that database size may skew results, especially when said databases are not capturing related information. For instance, including the immunologic signatures collection (c7) from the MSigDB in a vissE analysis of non-lymphoid cancer cell lines will bias some of the results towards immunologic phenotypes because of the large size of this collection (5219 gene sets in v7.2), despite these cell lines not having an immune phenotype.

This is a specific instance of a more general limitation that applies to gene-set enrichment analysis: biological processes and phenomena that are widely studied will be overrepresented in knowledgebases and will therefore skew results of enrichment analysis. Due to these limitations, it is important to choose related knowledgebases when performing a vissE analysis. Our recommendation for studying cancer systems and other non-lymphoid systems is to use the hallmark collection (h), the canonical pathways sub-collection (CP) of the curated gene-sets collection (c2), the cell type signatures collection (c8) and the ontology collection (c5) excluding the human phenotype ontology (HPO) of MSigDB. Other subcollections should be included in a study-specific manner. Similarly, vissE and other summarisation tools inherit limitations of gene-set enrichment analysis. Importantly, since this is a knowledge-driven tool, the discoveries made using vissE will be limited to known pathways and biological processes. However, vissE does allow exploration of the relatedness of processes in the biological system being studied, supporting the discovery of context-specific phenotypes. Though unknown processes cannot be identified, their presence can be suggested by vissE due to a guilt-by-association: the unknown process is likely to interact with other known processes and the vissE graph can show how these known processes are associated, leading to plausible hypothesis and potential explanations regarding the unknown process.

An important analytical module in the vissE arsenal is the text-mining analysis of gene-set clusters that facilitates cluster interpretation. The results of this analytical module, like any other analysis tool, depend on the quality of the underlying data. Concisely named gene-sets accompanied with succinct short descriptions would result in informative and interpretable word clouds. Curated knowledgebases such as pathway databases and GO generally use a controlled vocabulary to represent biological processes and are therefore rich information sources for text mining. The results in this study primarily used these sources and the resultant word clouds were biologically meaningful and easy to interpret. Consistent word clouds from text mining of names and short descriptions (Additional File 1: Supplementary Figure 1) attested to this claim and motivate our selection of specific sub-collections from the MSigDB. Collections such as the chemical and genetic perturbations (CGN) in the MSigDB contain many informative gene-sets however these have been named by individual contributors without a consistent naming convention and are limited in their utility in a text mining analysis. Concise, functional naming of gene-sets in repositories provides valuable information for downstream analysis of results and should be encouraged by knowledgebases.

Combining factor analysis with gene-set enrichment analysis and vissE, we were able to demonstrate a novel pipeline for unsupervised identification and characterisation of molecular phenotypes in various data modalities. Factor analysis has been previously used to explore expression patterns in an unbiased way however, the extension of this pipeline with singscore and vissE allowed us to gain a multifaceted view of the phenotype underlying the factors identified. Through this pipeline, we were able to identify and characterise proliferating cells in single cell transcriptomic data and the more nuanced phenotype of tumour promoting cancer associated fibroblasts (CAFs) in spatially resolved transcriptomics data. Despite capturing linear relationships in the data, the factor analysis algorithm we used proved to be powerful when combined with a functional interpretation pipeline. Since factors identified using PCA are orthogonal, we expect that biological processes captured using it are also orthogonal. The same process appearing across different factors would likely represent different context-specific states that produce context-specific outcomes. We expect PCA to perform better than other sophisticated approaches because it can capture modules of co-occurring context-specific processes within orthogonal factors that can then be decoupled using vissE. Other approaches such as independent components analysis (ICA) may reveal independent processes that biologists would then have to investigate for associations. The choice of PCA was easily justified with the results of the spatial transcriptomics analysis: using our pipeline, we were able to recover and characterise spatial structures associated with complex molecular phenotypes despite not having used the spatial context in the analysis. These results showed that spatially resolved transcriptomic data has the potential to recapitulate fine-grained spatial structures using purely transcriptomic measurements. The problem then becomes associating these gene expression signatures with known biology, which is in essence the problem that vissE has been designed to address.

The tool presented here, vissE, takes us a step forward in gaining a more holistic view of biological systems when coupled with state-of-the-art statistical methodology, and importantly, helps to remove investigator bias in interpretation.

## Declarations

### Availability of data and material

The vissE method along with gene-sets from the MSigDB can be accessed using the R/Bioconductor packages `vissE` and `msigdb`, respectively. Data from the EMT study can be accessed using the `emtdata` R/Bioconductor package. Input data to run vissE on the three workflows in this study are available on figshare (DOI: 10.26188/19193660).

### Supplementary data

Additional file 1: Additional figures to support the analyses in this study.

Additional file 2: Top 60 biological themes identified from the vissE analysis of the EMT system.

Additional file 3: Top 60 biological themes identified from the vissE analysis of the fourth principal component of the single-cell transcriptomic data from 6 breast cancer patients.

Additional file 4: Top 60 biological themes identified from the vissE analysis of the first principal component of the 10x Visium spatial transcriptomic data from a breast cancer patient.

### Authors’ contributions

DDB and MJD conceived and led the study. DDB, NL, SL, MK and AM performed the computational and statistical analysis. DDB, MK and CWT developed the R/Bioconductor packages used in this study. DDB, CWT, HJW and MJD assisted in the interpretation of results. NP annotated the pathology slides from the spatial analysis. MJD supervised the study. All authors helped in preparing the manuscript.

