## Supplementary figures and images for "vissE: A versatile tool to identify and visualise higher-order molecular phenotypes from functional enrichment analysis"

### Additional File 2

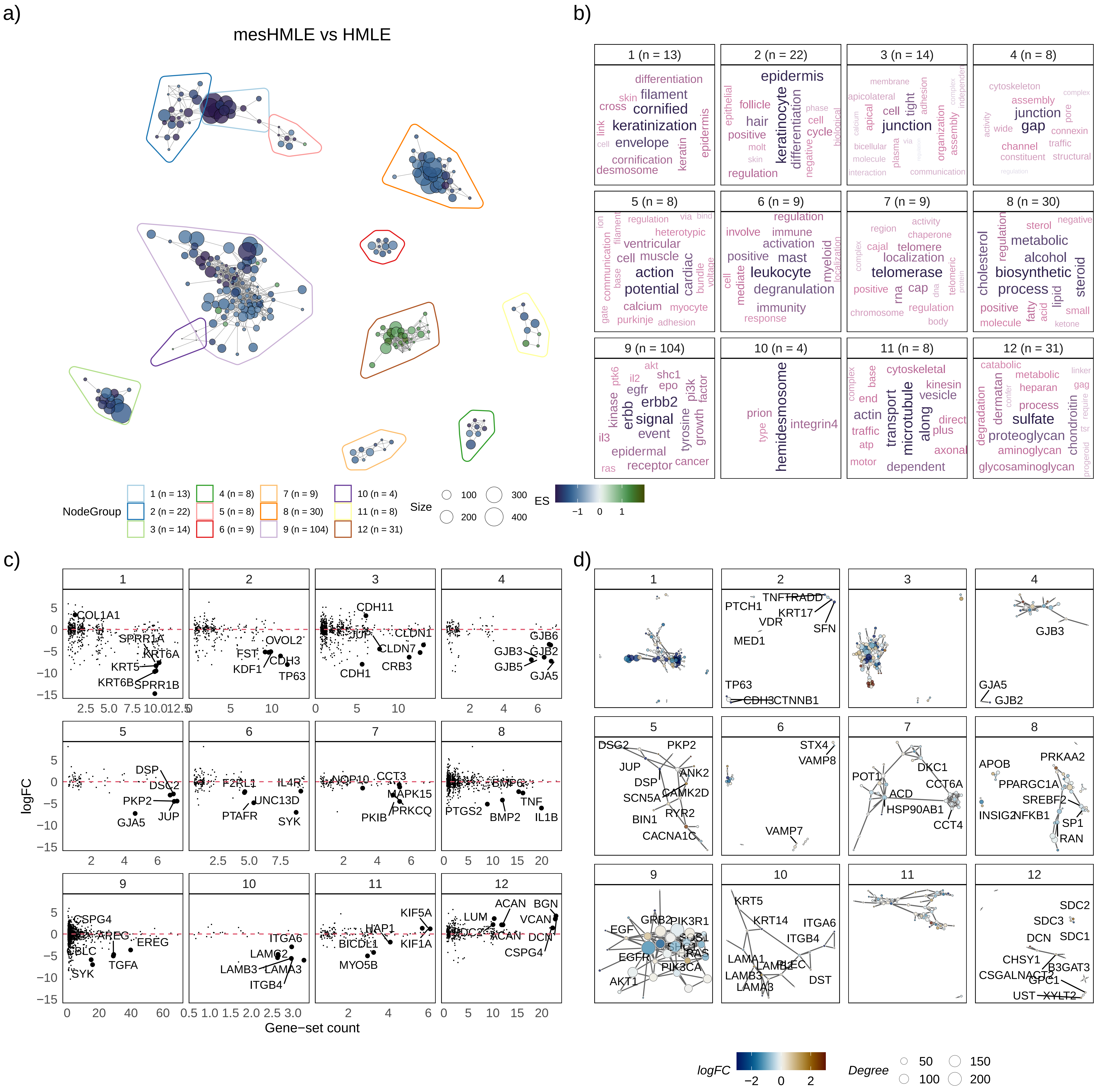

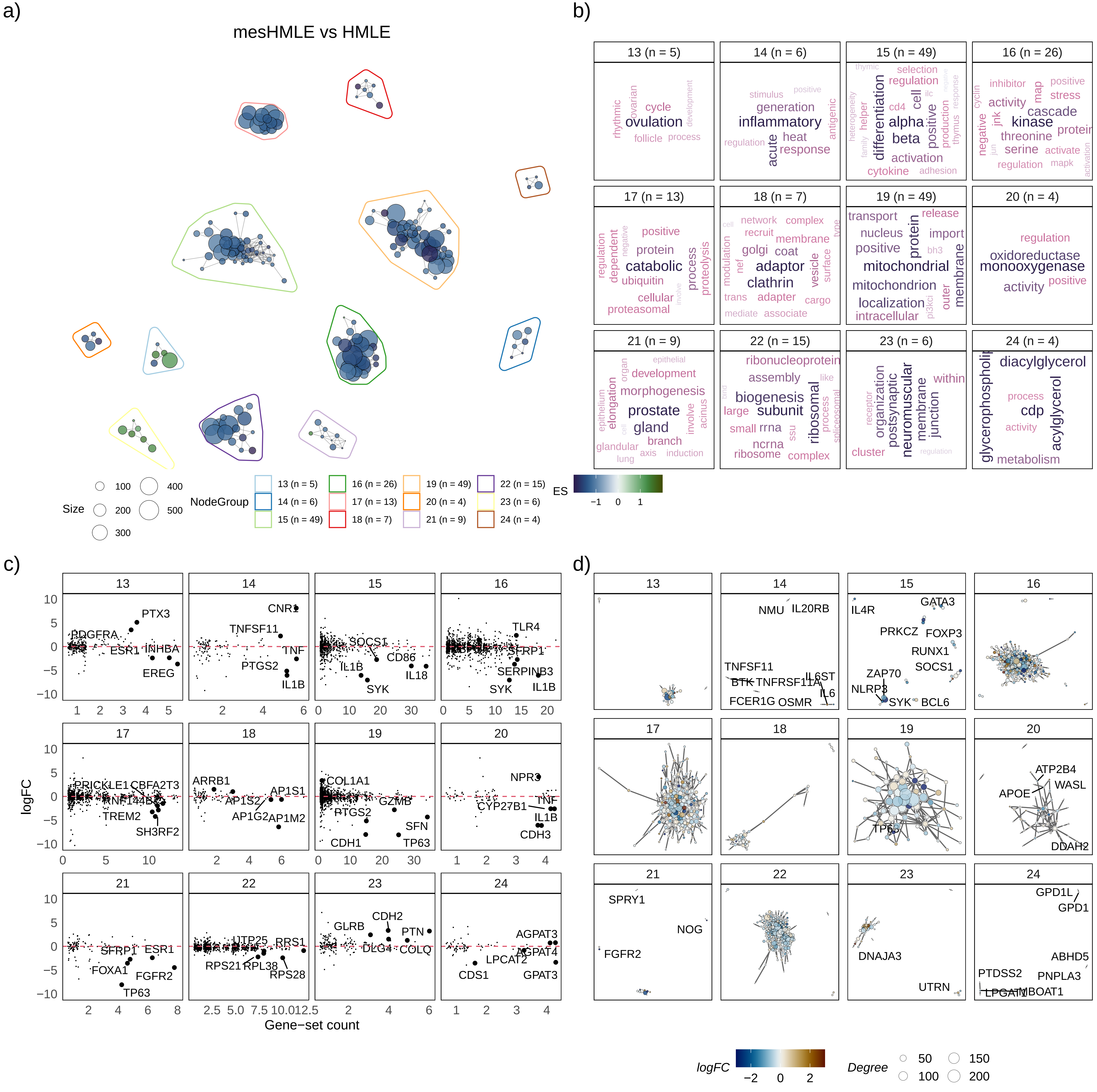

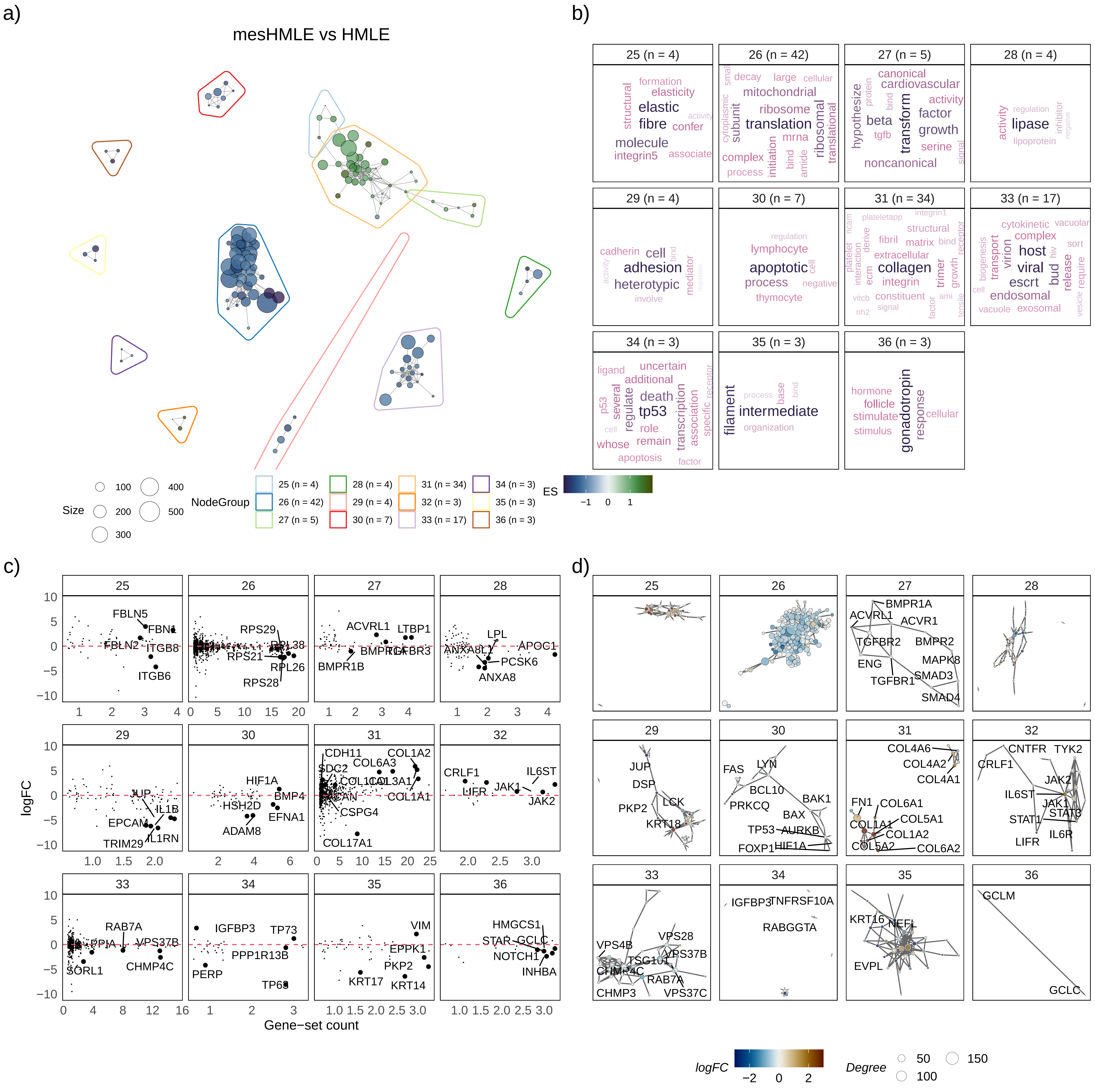

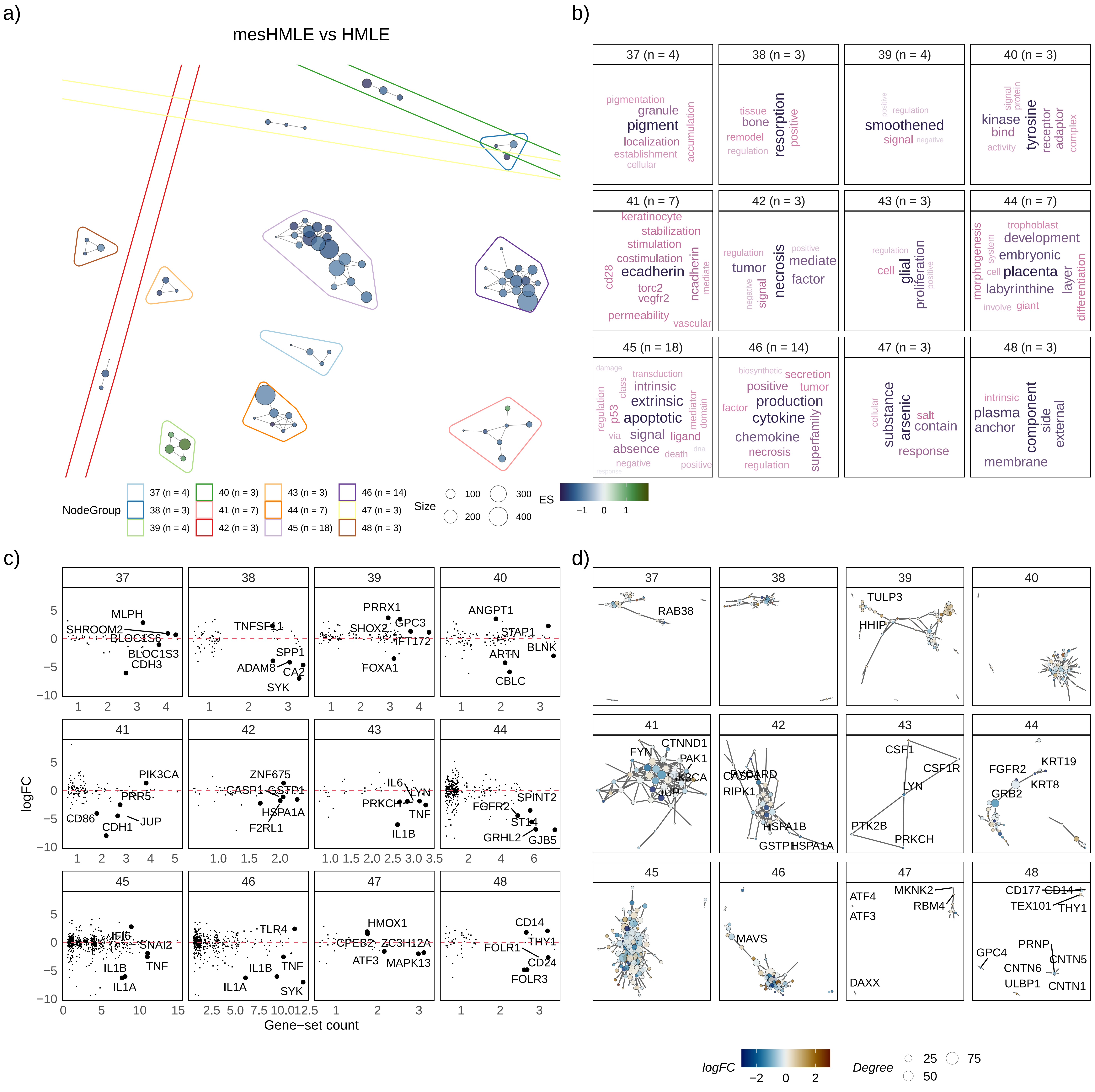

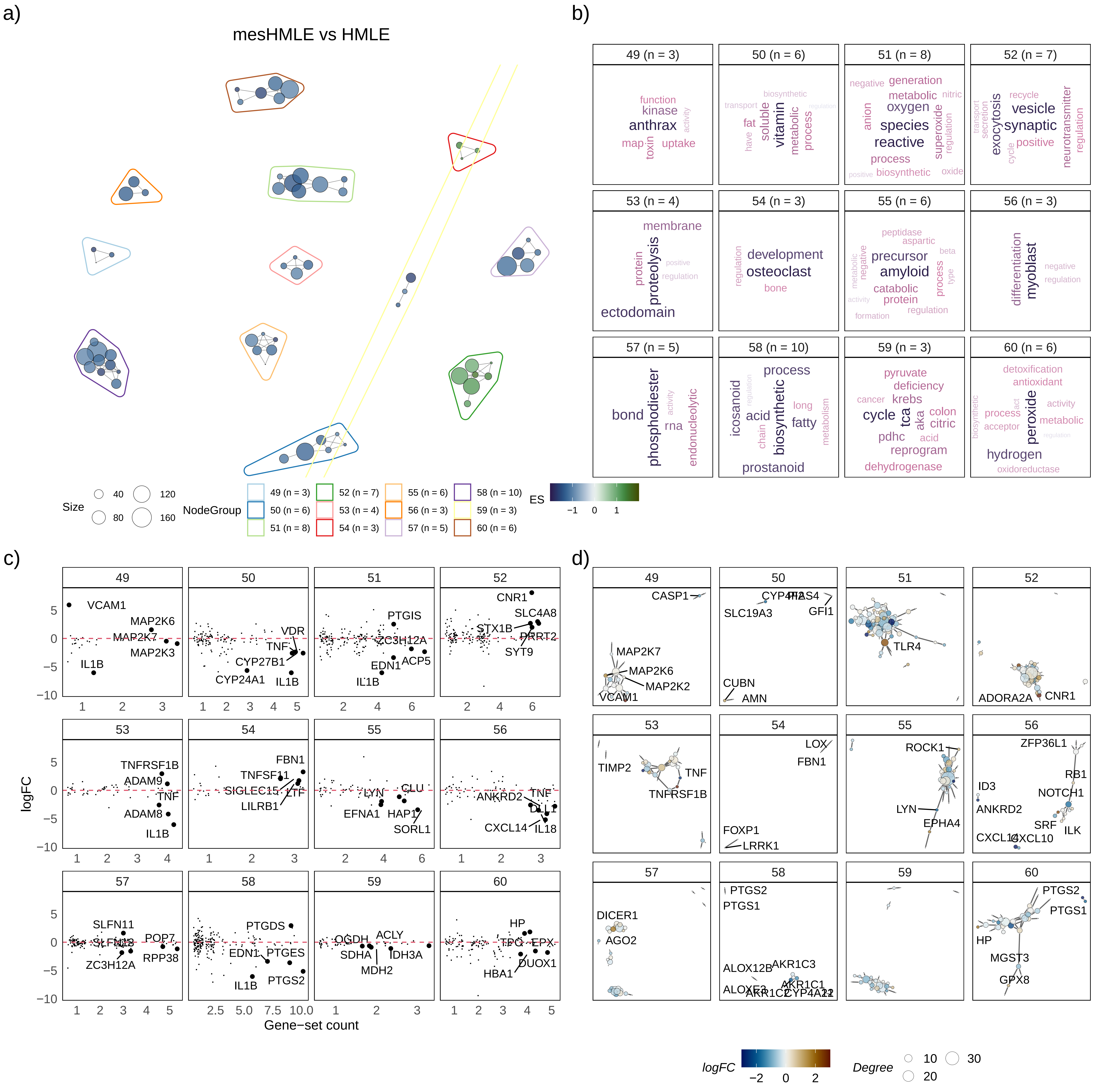

### Additional File 3

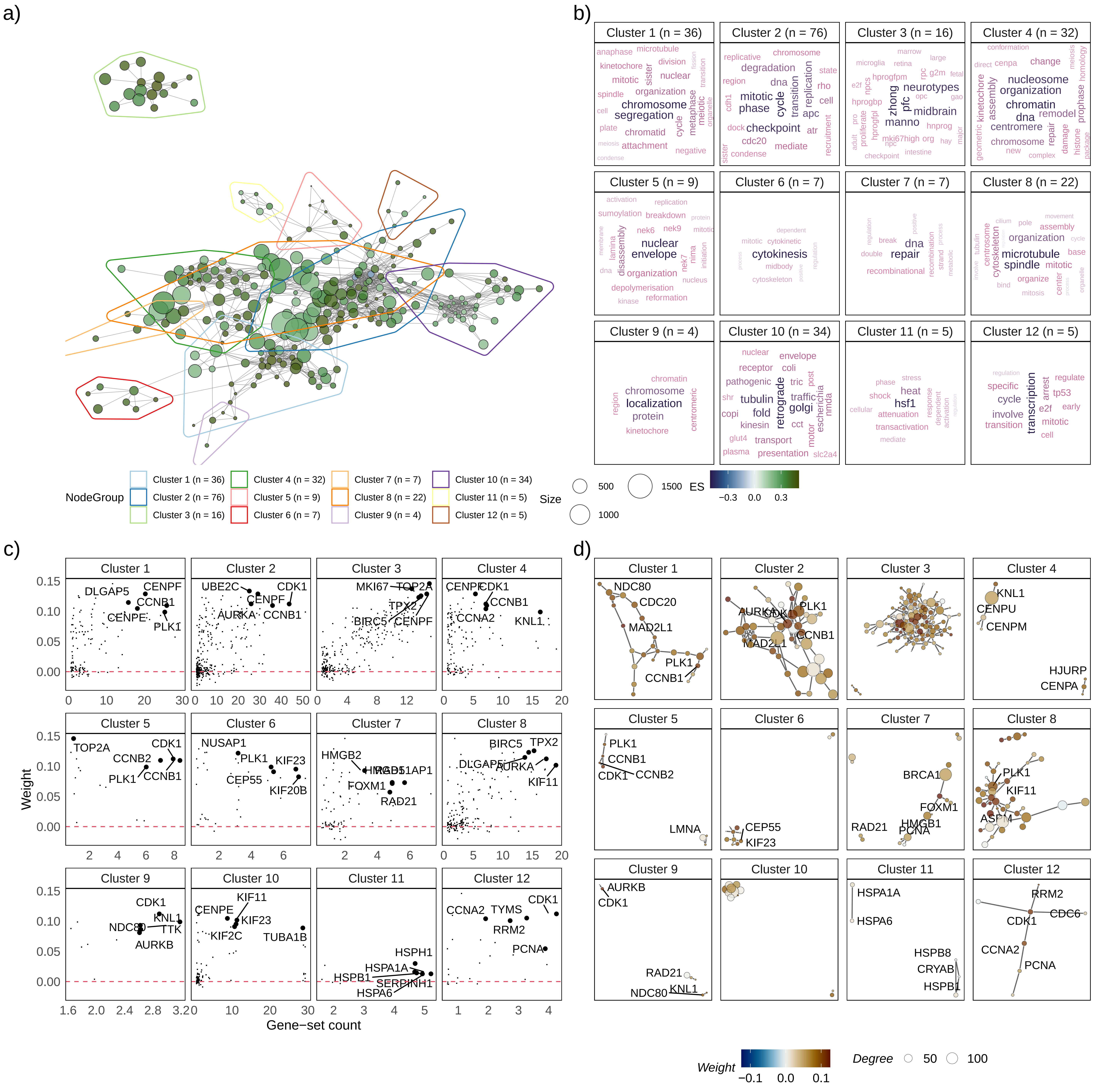

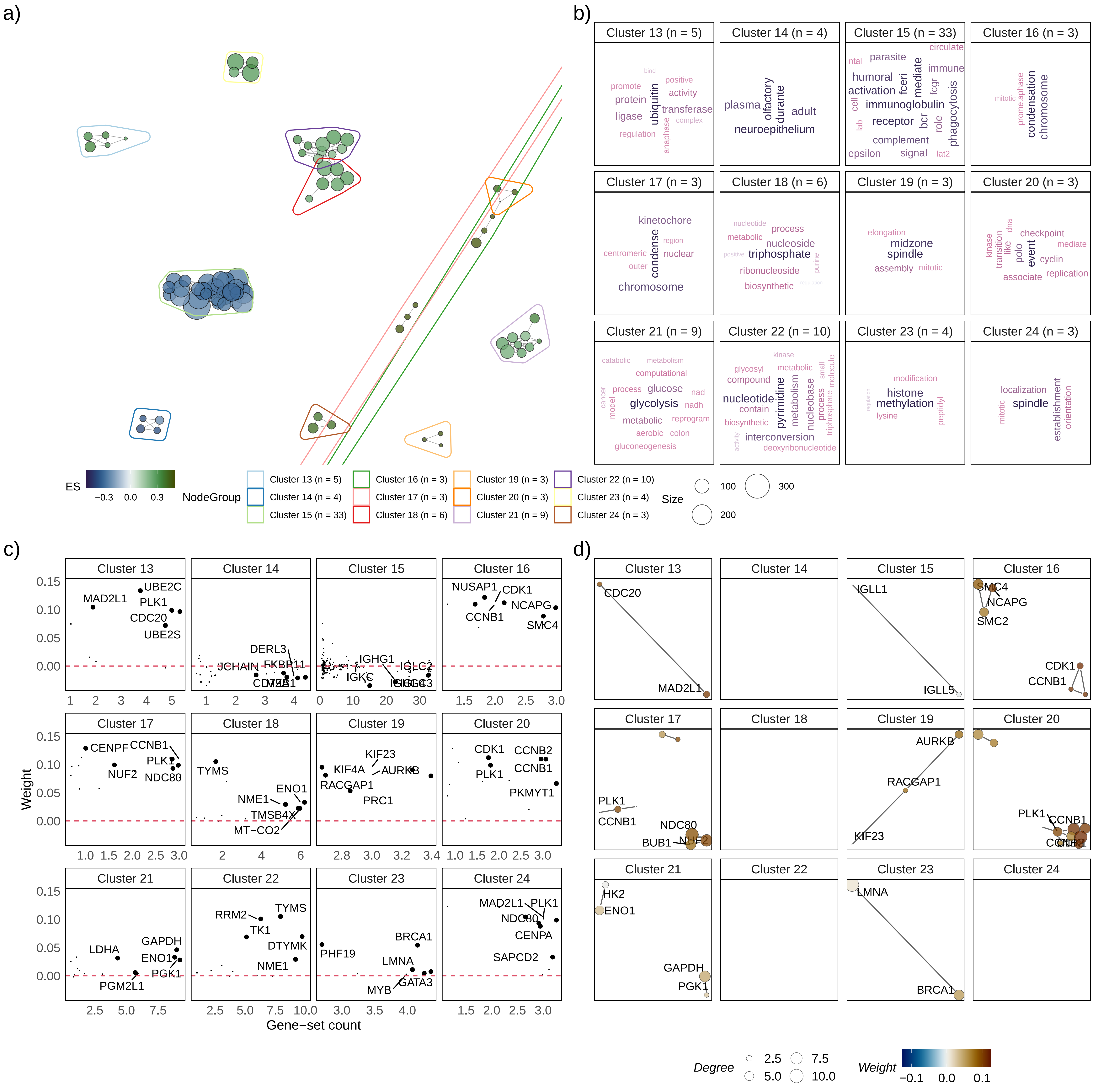

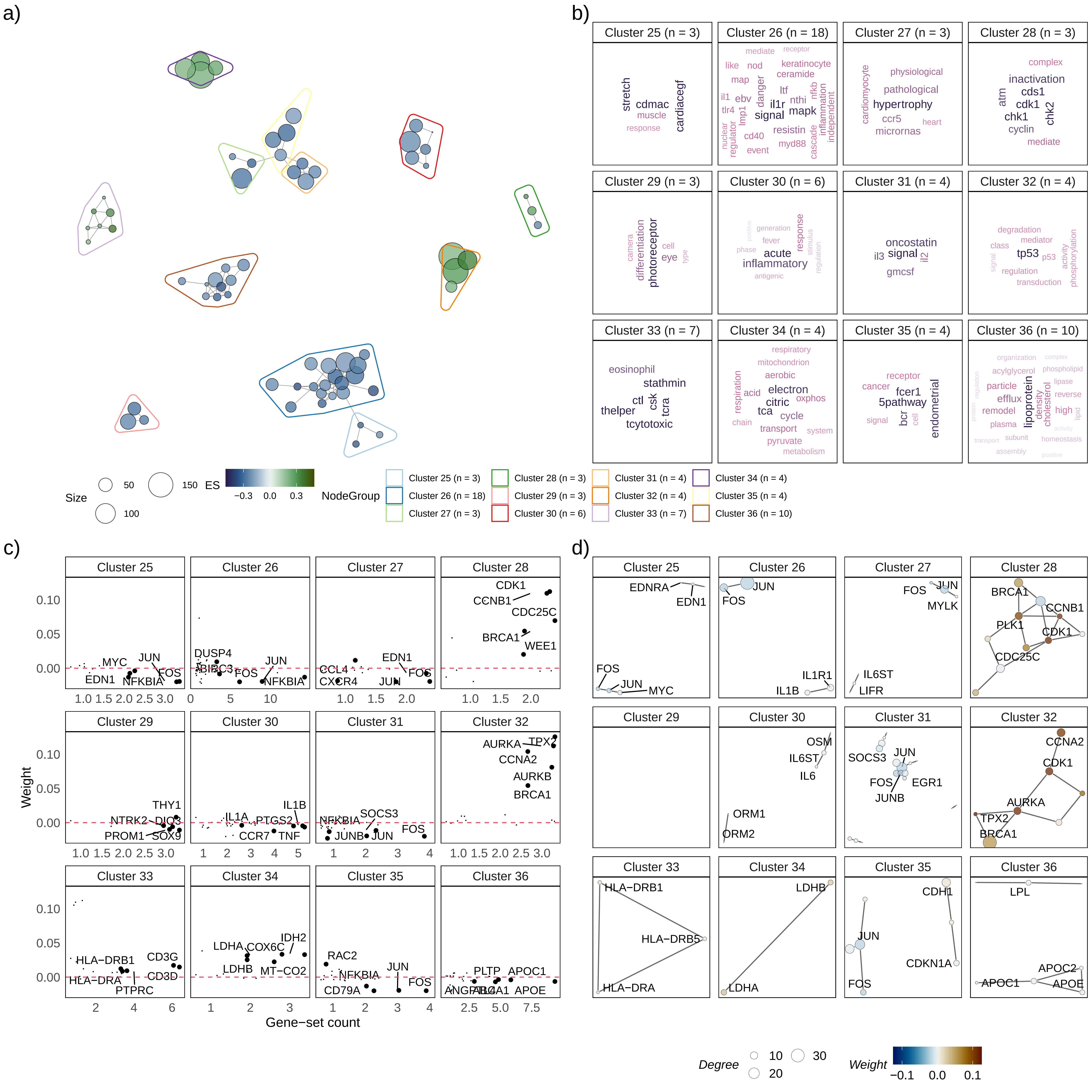

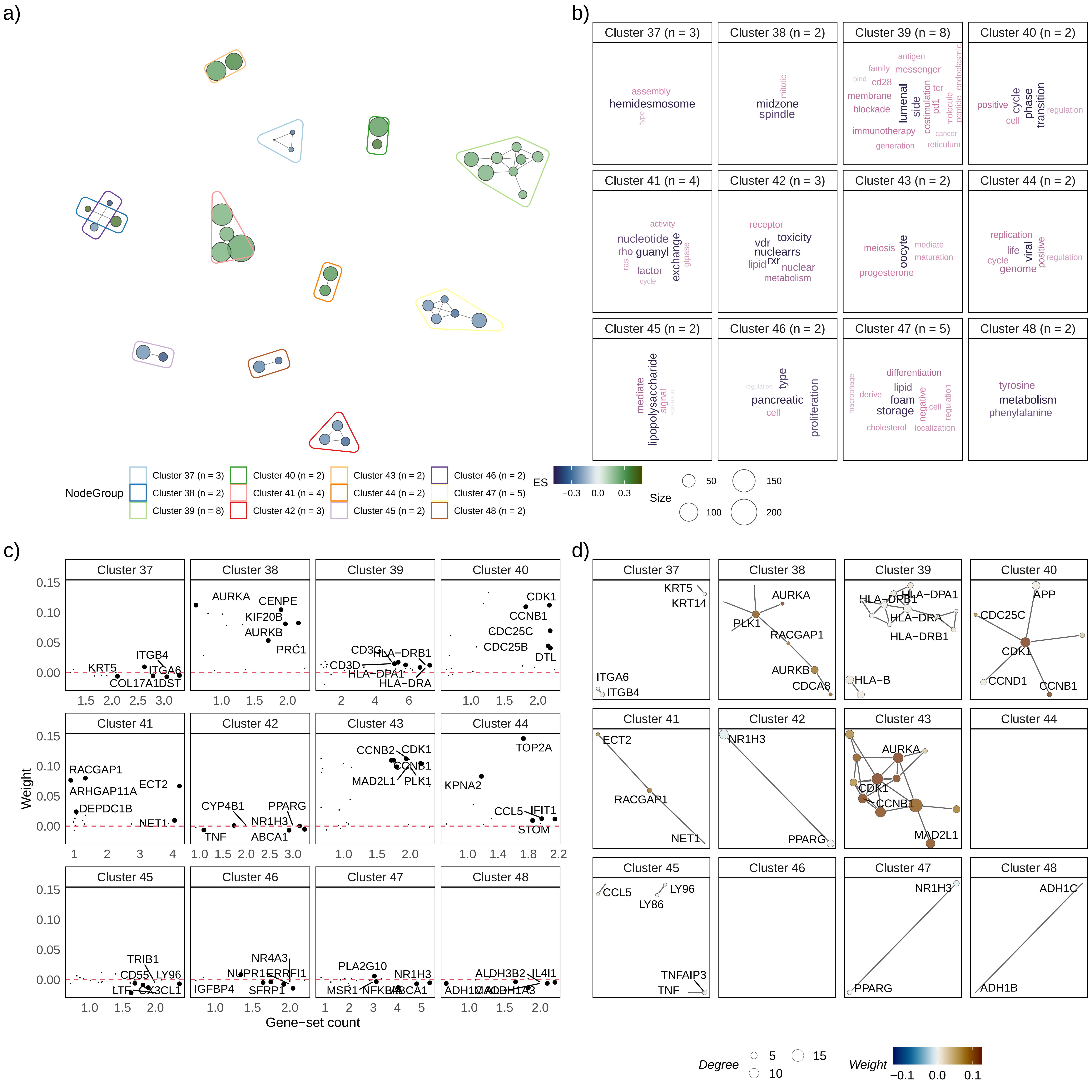

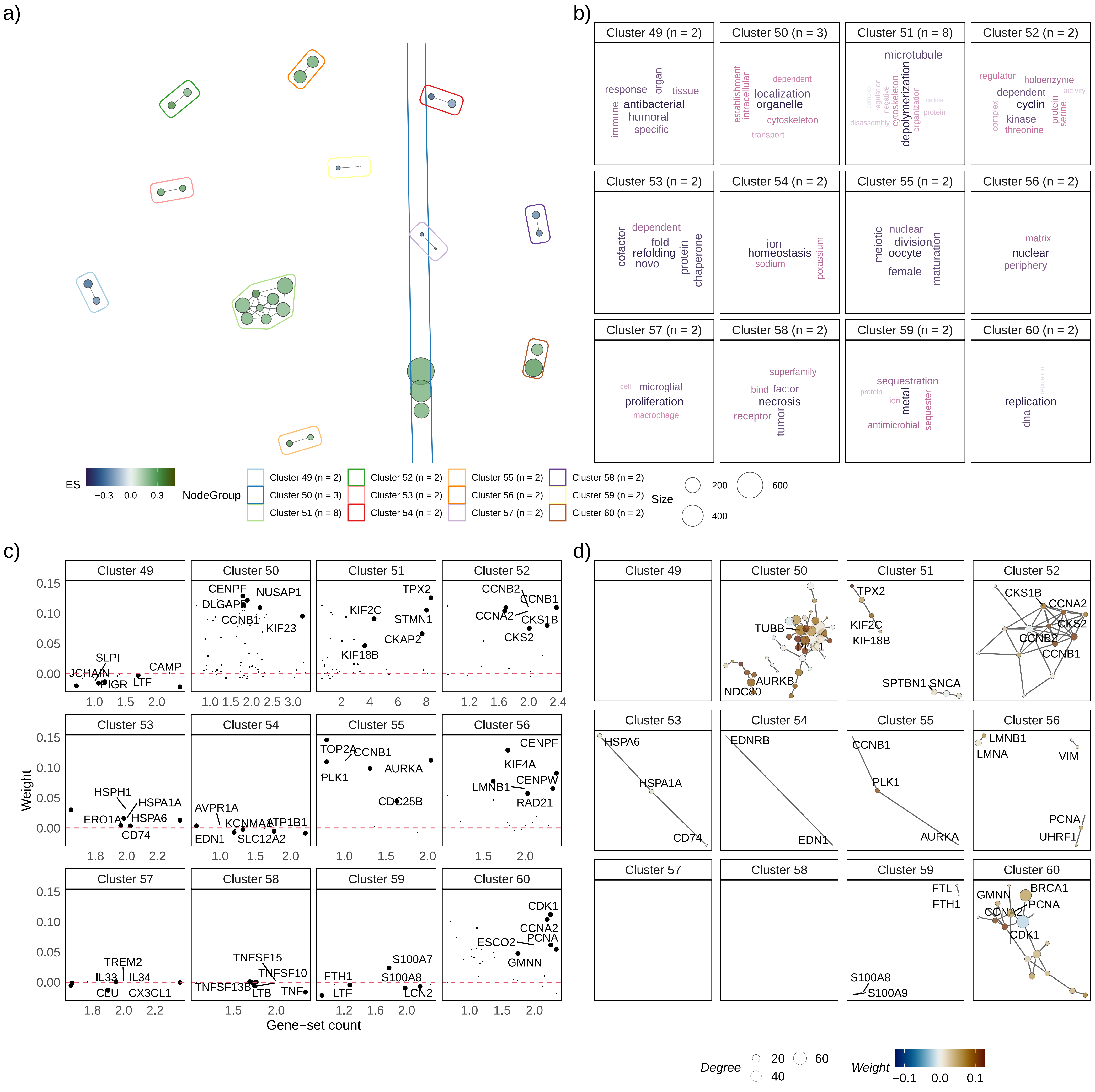

### Additional File 4

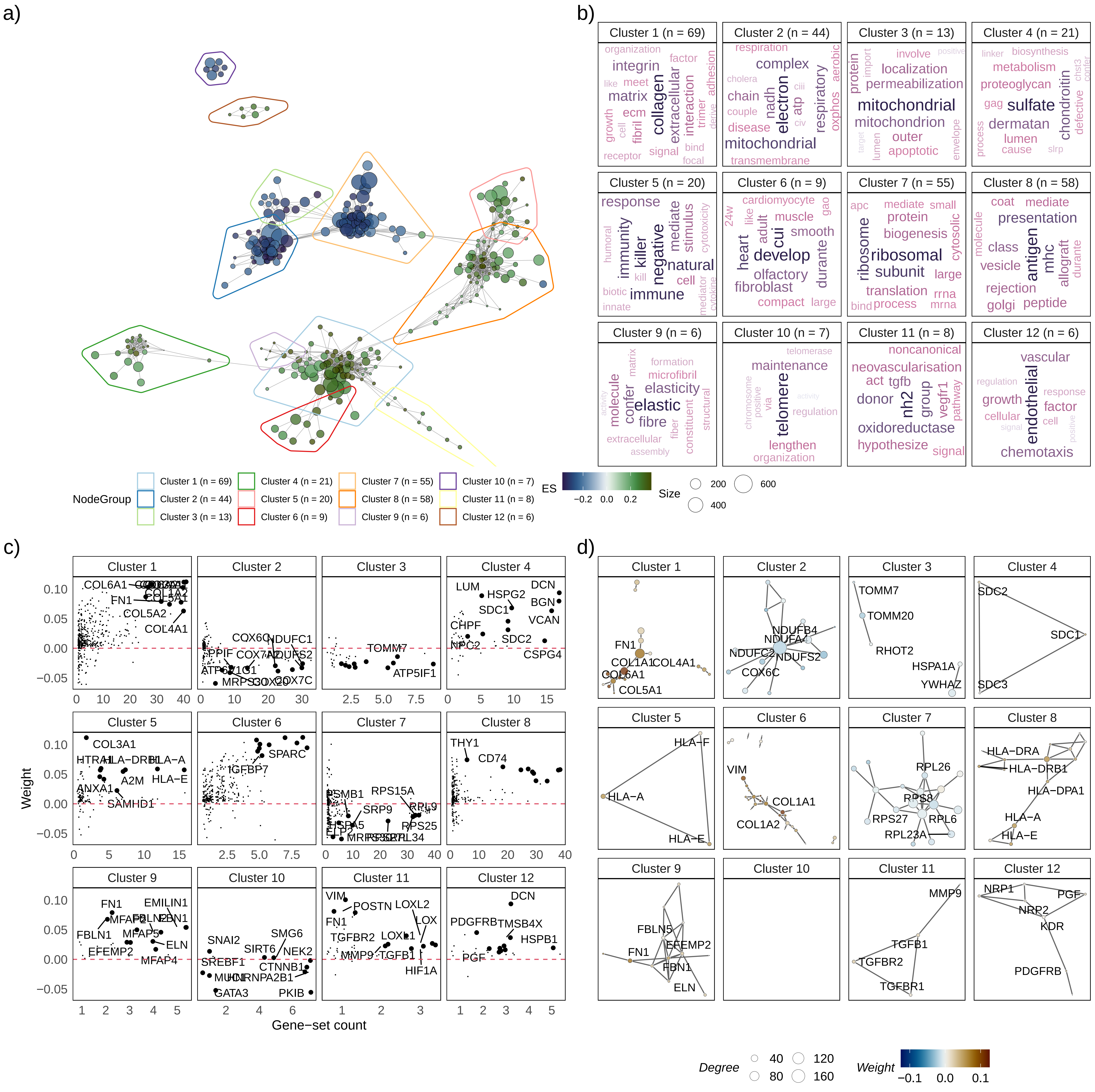

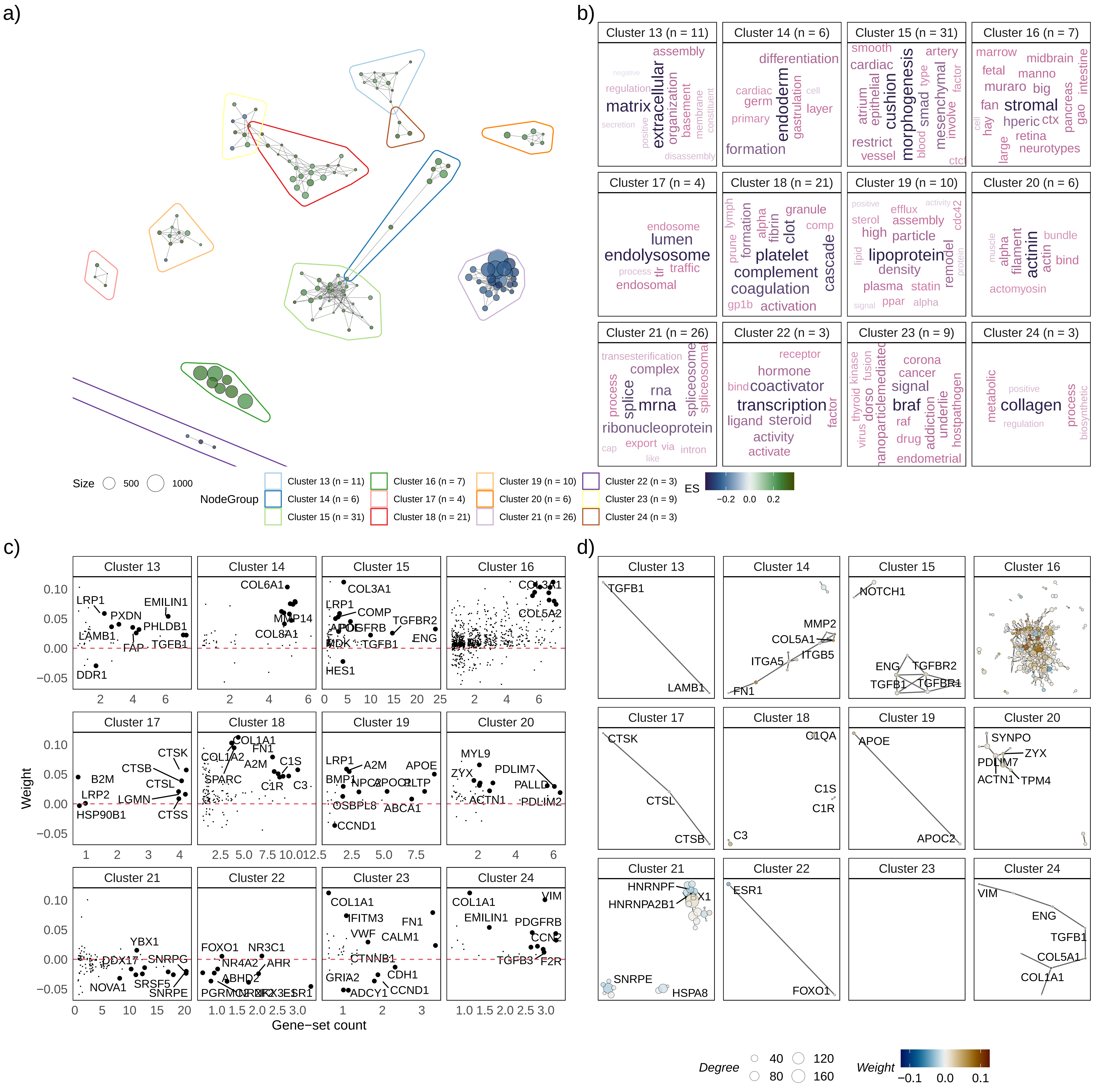

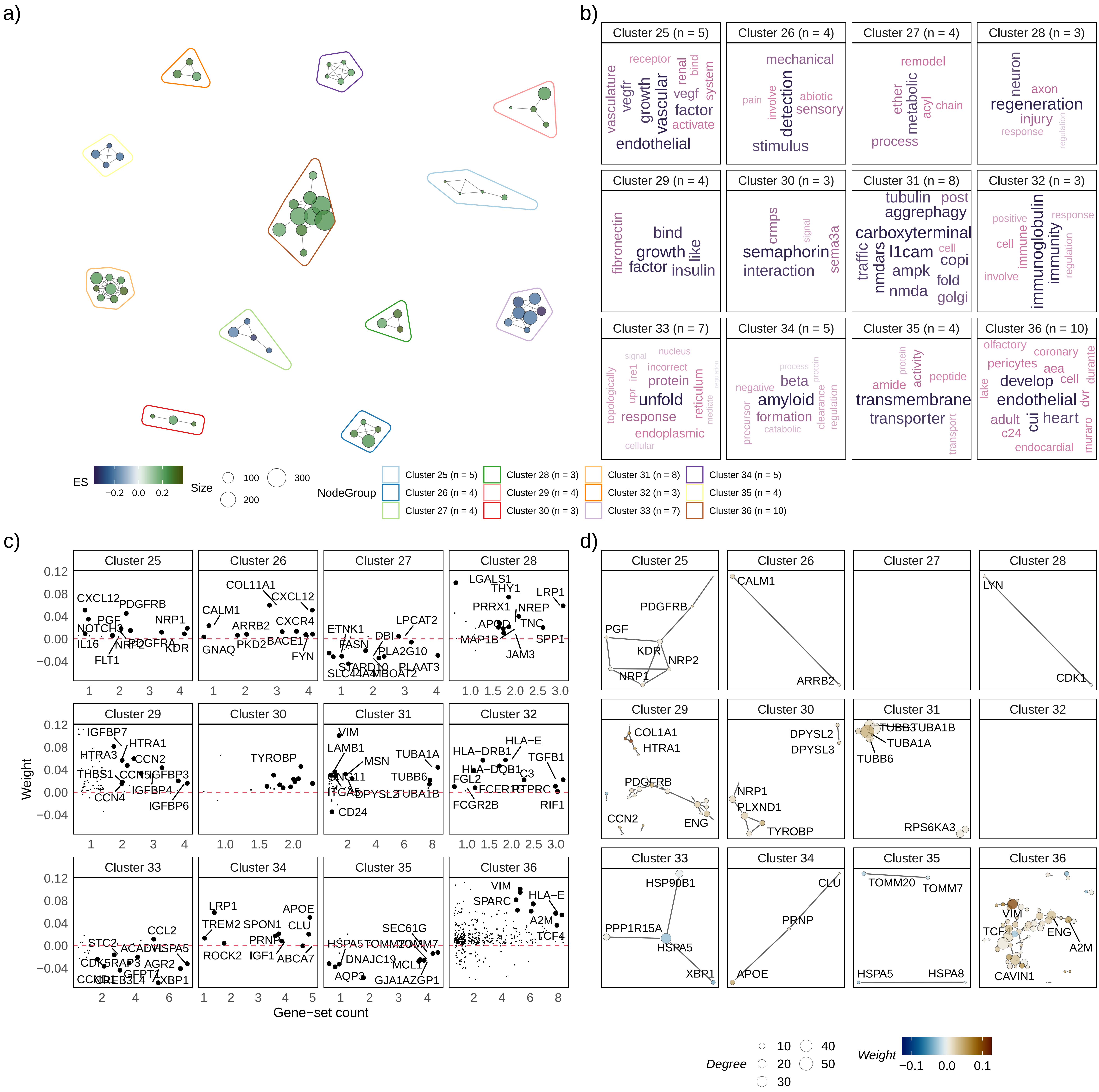

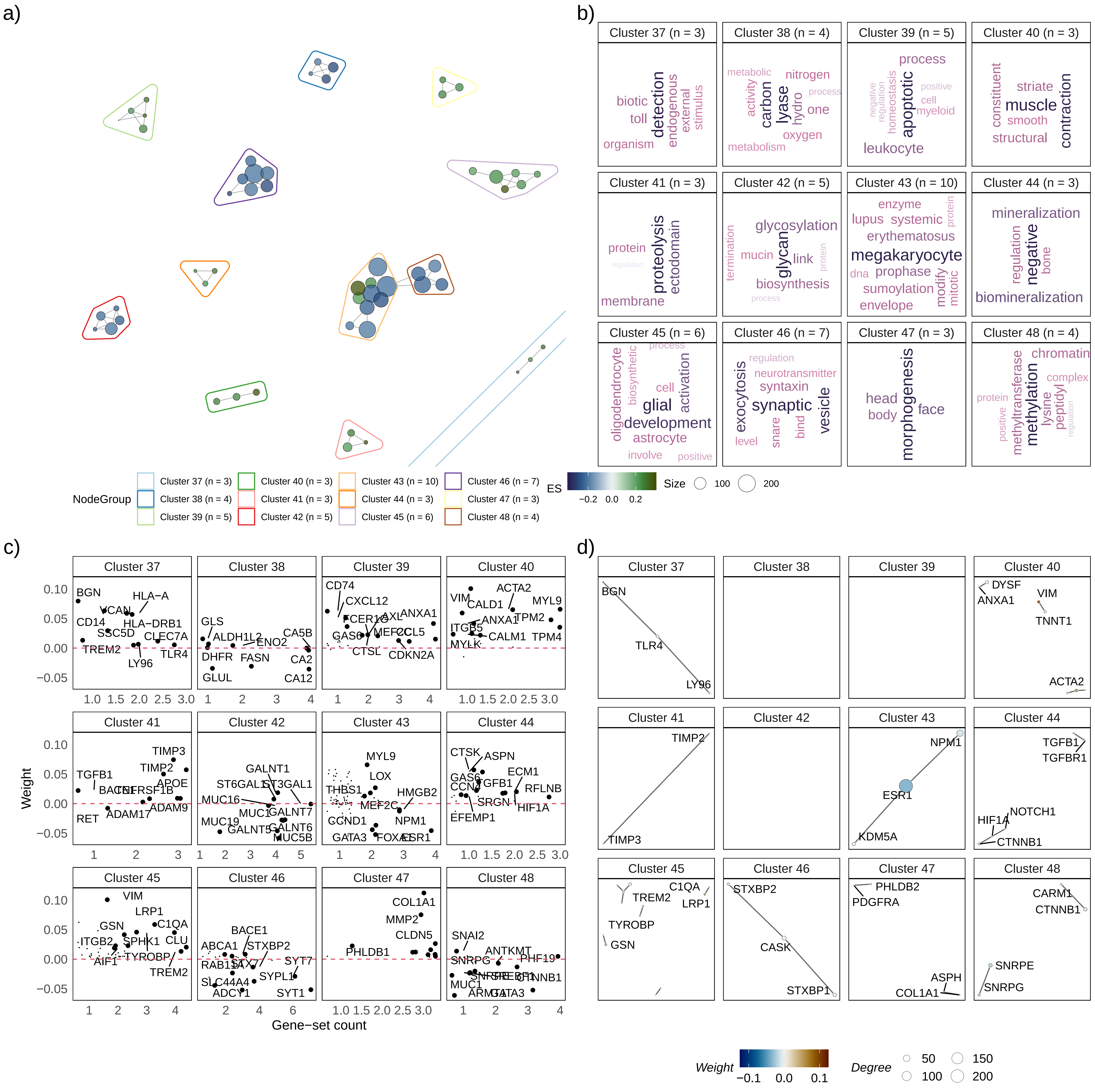

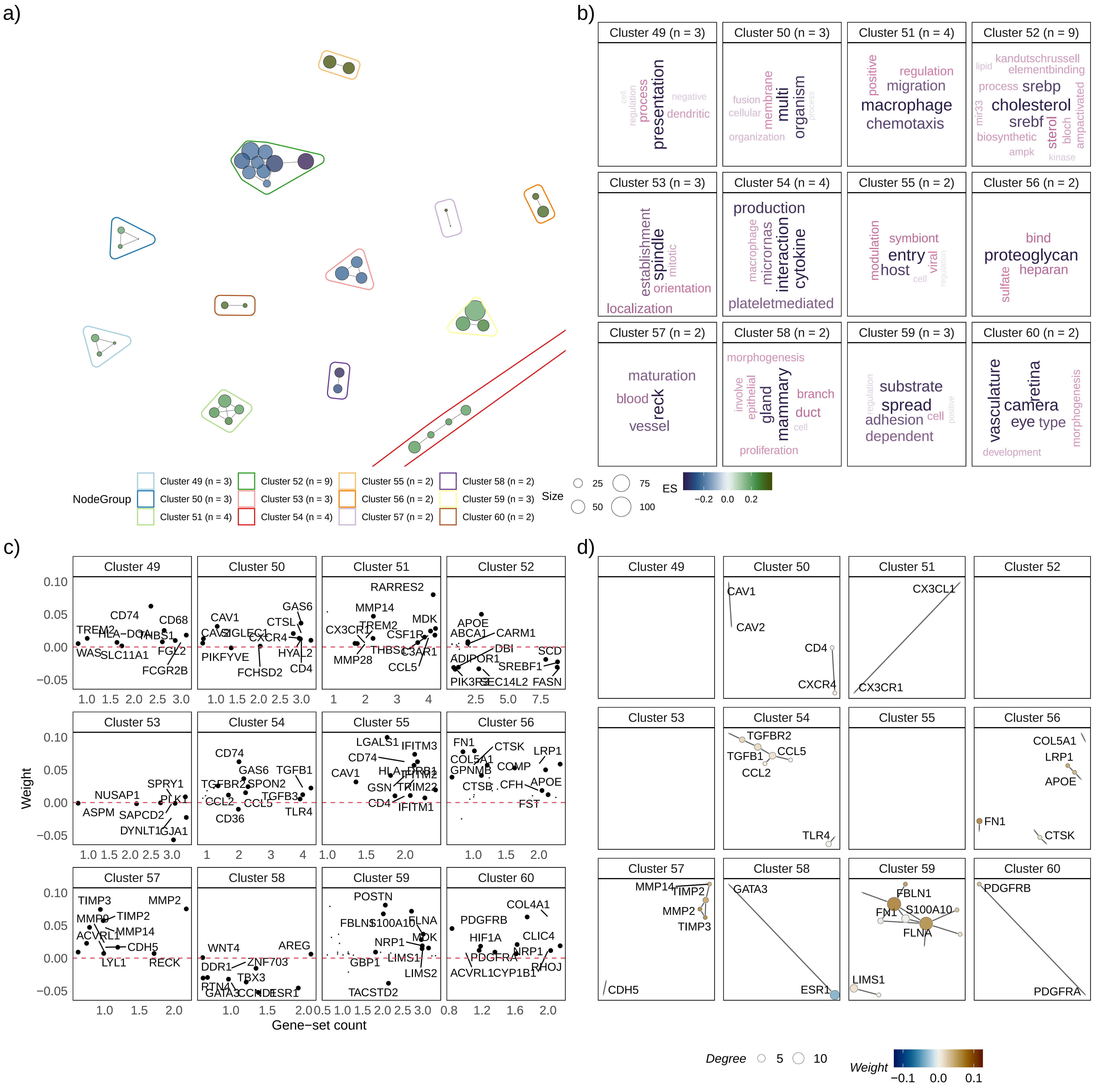
