## Additional File 1 for "vissE: A versatile tool to identify and visualise higher-order molecular phenotypes from functional enrichment analysis"

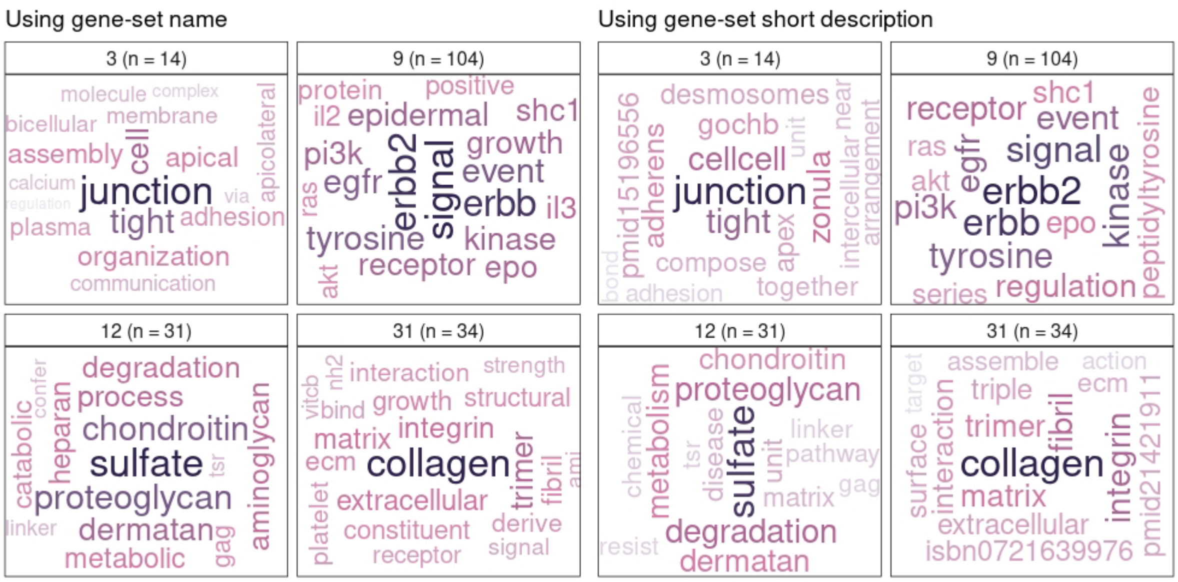


**Supplementary Figure 1:** The same biological themes are inferred whether using gene-set names or their short descriptions to annotate gene-set clusters identified using vissE.

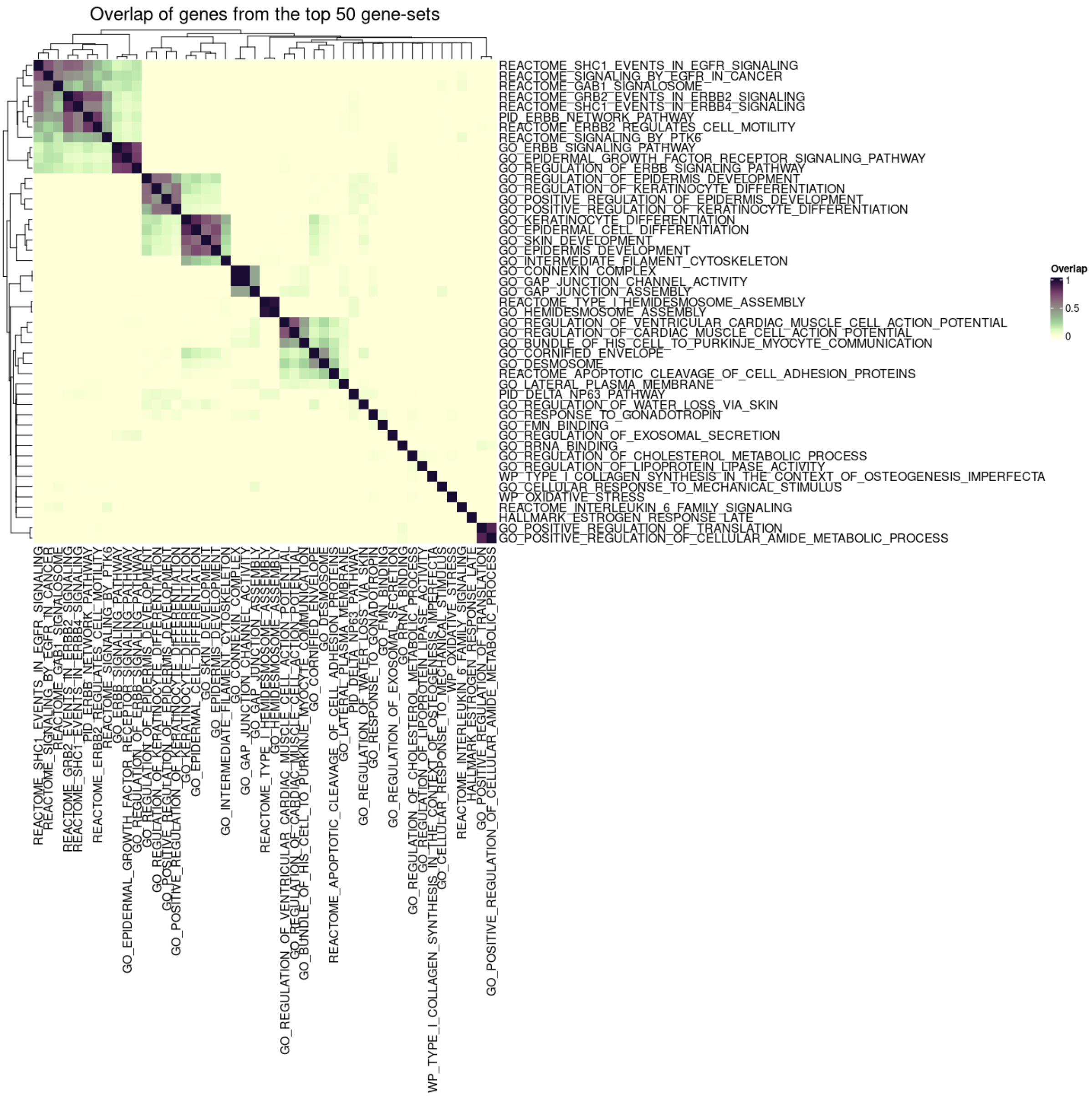


**Supplementary Figure 2:** The top 50 significant gene-sets in the HMLE system differential expression analysis shared a large number of differentially expressed genes and thus were representing redundant functional information.

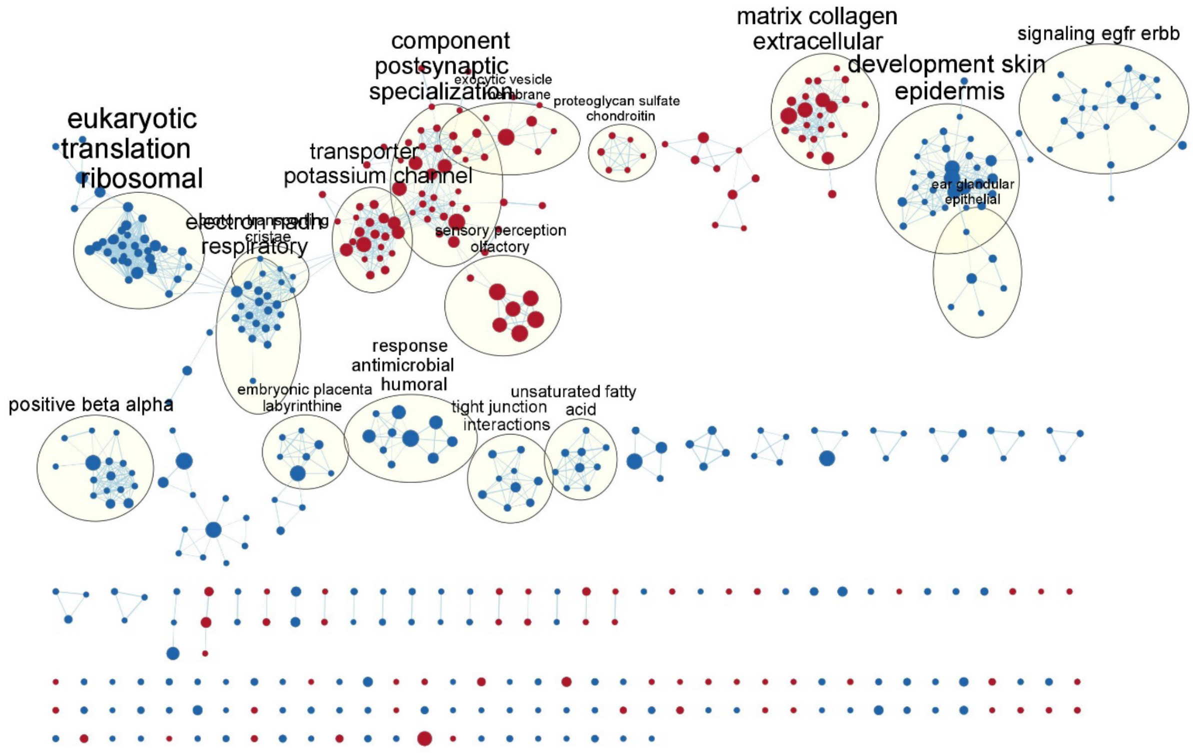


**Supplementary Figure 3:** Gene-set clusters and their annotations derived from EnrichmentMap to characterise the epithelial to mesenchymal transition in the HMLE system. Each node represents a gene-set and edges connect similar gene-sets. Nodes are coloured based on the direction of change: red – upregulated in mesenchymal cells; blue – upregulated in epithelial cells.

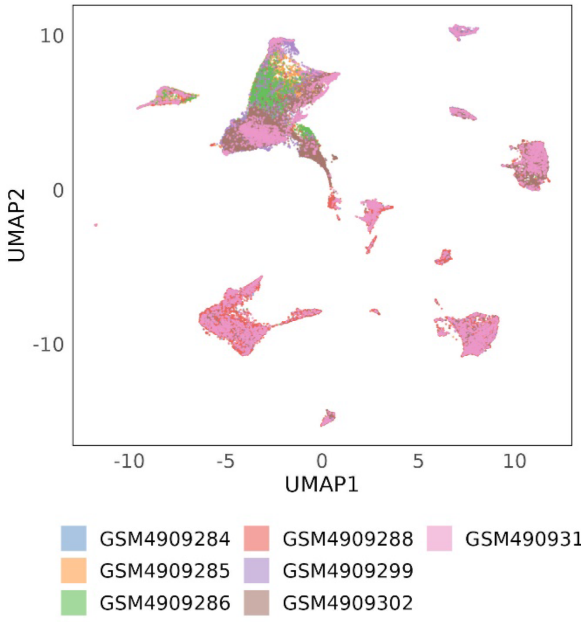


**Supplementary Figure 4:** UMAP of cells from seven breast cancer patients representing two breast cancer subtypes. Cells are coloured based on the patient they were derived from.

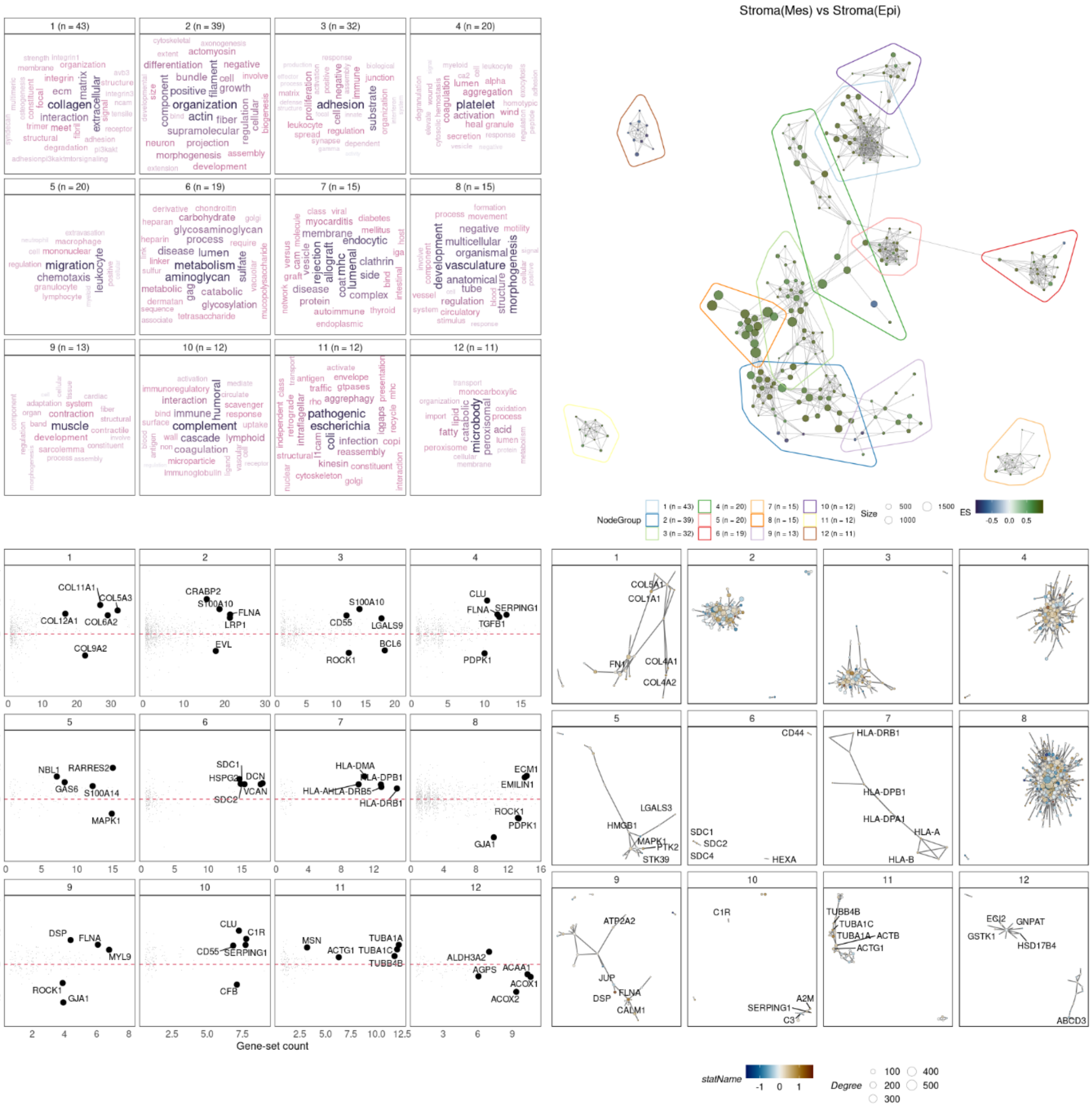

**Supplementary Figure 5:** The top 12 themes identified from a vissE analysis of the results of a differential expression analysis between stromal spots surrounded by malignant epithelial cells vs. stromal spots surrounded by malignant mesenchymal cells.
